# Wheat infection by *Fusarium graminearum* species complex members is facilitated by a transcriptionally conserved non-ribosomal peptide synthetase gene cluster

**DOI:** 10.1101/2025.01.24.634825

**Authors:** Sabina Moser Tralamazza, Mikkel Rank Nielsen, Leen Nanchira Abraham, Anna K. Atanasoff-Kardjalieff, Gaetan Glauser, Jens Laurids Sørensen, Lena Studt-Reinhold, Benedito Corrêa, Daniel Croll

## Abstract

Biosynthetic gene clusters (BGCs) are often found in fungal pathogens and encode secondary (or specialized) metabolites that play crucial roles in host and niche adaptation. However, the regulatory dynamics and functions of these BGCs during host infection remain largely unknown. To address this gap, we employed interspecies comparative transcriptomics to identify BGCs involved in host-pathogen interactions mediated by the *Fusarium graminearum* species complex (FGSC). We conducted joint transcriptomic and metabolomic analyses during wheat infection and *in vitro* experiments with five members of the FGSC to understand gene regulation during host infection. Our findings revealed that expression regulation was predominantly species-specific, but we also identified a set of shared upregulated genes that were common to all five FGSC species showing enrichment for metabolic and pathogenicity-associated functions. We focused on jointly upregulated BGC during infection and identified a NRPS-like gene cluster named SM3. The cluster was highly conserved within FGSC and shared by the more distant *F. sambucinum* species complex. We show that inactivation of the BGC core gene significantly impairs *F. graminearum* sensu stricto (s.s.) spore production, reduces fungal spread and Fusarium Head Blight symptoms on wheat kernels. Our results highlight that comparative infection transcriptomes can reveal conserved metabolic functions mediating the host infection process of a plant pathogen.

## INTRODUCTION

Secondary or specialized metabolites (SM) play a pivotal role in mediating interactions between fungal pathogens and their host, fueling an evolutionary “arms race” (Maor & Shirasu, 2005), yet most SM functions remain largely unknown (Rokas et al., 2020). SMs typically encode metabolic pathways that are dispensable for basic cellular survival but provide accessory characteristics enabling organisms to more effectively adapt to ecological pressures (Rokas et al., 2018), promote defense mechanisms (Cordero and Casadevall 2017; Won et al. 2022), secure nutrition (Harrison et al., 2022) and promote virulence (Kessler et al., 2020; Patron et al., 2007; Proctor et al., 2018; Yamada et al., 2000). For example, HC-toxin is a cyclic tetra-peptide that functions as a virulence factor for the fungus *Cochliobolus carbonum* during maize infection (Brosch et al. 1995). Similarly, deletion of the metabolites botcinic acid and botrydial significant hampers infection by *Botrytis cinerea* in tomato crops (Deighton et al., 2001). Genes involved in SM biosynthesis are generally clustered as a biosynthetic gene cluster (BGC) with each gene encoding a specialized biosynthetic function (Nützmann et al., 2018; Rokas et al., 2020). BGCs encode pathways to produce primarily four main chemical classes: polyketides (PKS), non-ribosomal peptides (NRPS), terpenes and indole alkaloids or hybrids thereof. The core biosynthetic gene encodes for the backbone chemical structure of the metabolite while tailoring genes with diverse enzymatic functions modify, regulate or transport the secreted metabolite outside the cell (Keller, 2019).

Phylogenetic conservation of metabolic processes facilitates the search for targets with relevant cellular functions (Ebert et al., 2019; Peregrín-Alvarez et al., 2009). Conserved BGCs often indicate critical metabolic processes (Arulprakasam & Dharumadurai, 2021; De Jonge et al., 2018; Klassen, 2010; Rank et al., 2011), that interconnect with the primary metabolism (García-Estrada et al., 2018; J. K. Hicks et al., 1997; Wilkinson et al., 2004) and play key roles in fungal growth, development and pathogenesis (Calvo et al., 2002). However, regulation and production of SMs vary significantly among species (Amos et al., 2017), which can pose challenges for effective metabolite discovery. The activation of BGCs is a multifaceted process influenced by a complex network of local and master modulators (Keller, 2019). These modulators respond to environmental cues such as temperature, light, pH, and nutrient availability (Cervini et al., 2021; Espeso & Peñalva, 1996; Won et al., 2022). Genetic variation and epigenetic mechanisms often impact the regulation of metabolic pathways (Atanasoff-Kardjalieff & Studt, 2022). Furthermore, species-specific and niche adaptations introduce an additional layer of complexity when studying SM activation under laboratory conditions (Boutigny et al., 2011; Drott et al., 2021; Harris et al., 2016; Takahashi et al., 2021). Highly conserved BGCs tend to be more expressed than less conserved clusters (Amos et al., 2017). Under such a scenario, conserved regulation across species during host infection could serve as an effective method for identifying relevant targets in pathogenic interactions (Amos et al., 2017). Such cross-species comparative transcriptomics has successful identified genes related with nitrate assimilation in fungi (J. C. Nielsen et al., 2019), phagosome maturation in plants (Bruno et al., 2021) and oxidative stress in bacteria (Giraud-Gatineau et al. 2024).

The *Fusarium graminearum* species complex (FGSC) comprises a major group of 17 fungal pathogen species affecting small cereals, including wheat, barley, and rice (van der Lee et al., 2015). They can cause a wide range of diseases, including seedling blight, crown rot, and Fusarium Head Blight (FHB) with *F. graminearum* s.s. the most prevalent species within the complex (van der Lee et al., 2015). Despite their genetic similarity (Wang et al., 2011), species exhibit remarkable diversity in terms of aggressiveness, host preference, and geographic distribution (de Chaves et al., 2022; Gomes et al., 2015; H. Zhang et al., 2012). Hence, species of the FGSC provide an ideal model system to investigate shared regulatory elements involved in the host infection process. FGSC is a major concern for global food safety due to their ability to produce mycotoxins, toxic SM in food and feed (Johns et al., 2022). Despite ongoing efforts, successful management of FGSC damage remains challenging. Fungicide resistance has steadily increased over the years (de Chaves et al., 2022), and some FGSC members remain insensitive to current control treatments (Yang et al., 2018). SMs play a crucial role in the pathogenicity of FGSC members. Deoxynivalenol, a member of the trichothecene mycotoxin group, acts as a major virulence factor following initial infection of wheat plants, facilitating the spread of the fungus within the rachis of wheat spikelets in *F. gramineaum* (Walter et al., 2010). Recently, the nonribosomal peptide fusaoctaxin A was identified as contributing to cell-to-cell invasion in *F. graminearum* s.s. (Jia et al., 2019). Some SM functions remain elusive including, for example, gibepyrone, a highly conserved type I PKS (PKS8) gene cluster across the *Fusarium* genus (Brown et al., 2022; Tralamazza et al., 2019). Evidence supports a moderate *in vitro* antimicrobial and parasitic activity (Barrero et al., 1993; Bogner et al., 2017; Janevska et al., 2016), yet the BGC’s contribution during infection remains poorly explored. Despite the importance of FGSC, research on pathogenicity factors remains predominantly focused on a single species, *F. graminearum* sensu strico (s. s.) (Brauer et al., 2020; Harris et al., 2016; Stephens et al., 2008; Tu et al., 2023). Furthermore, the majority of SMs encoded among FGSC genomes remain without known functions (Brown & Proctor, 2016; Sieber et al., 2014; Tralamazza et al., 2019).

Here, we used comparative transcriptomics in standardized environments across five FGSC members to determine the degree of co-regulation of SMs during host infection. We then focused on co-regulated BGCs across species as a strategy to uncover most likely candidates for BGCs associated with disease development. We tested the impact of the metabolite gibepyrone (PKS8/SM20) on virulence of *F. cortaderiae* and found no evidence of any infection impairment. Our approach led to the discovery of a NRPS BGCs named SM3. The BGC deletion significantly impaired *F. graminearum* s.s. spore production, markedly reduced fungal spread and Fusarium Head Blight symptoms in wheat kernels.

## METHODS

### FGSC genome annotation

We performed comparative genomics analyses with 16 FGSC species and three sister species belonging to the *Fusarium sambucinum* species complex (Supplementary Table S1). We annotated the genomes of the FGSC species, except for the previously annotated *F. graminearum* s.s. (PH1 - NRRL 31084) (King et al., 2017), *F. meridionale* (Fmer152), *F. cortaderiae* (Fcor30FRS), *F. austroamericanum* (Faus154) and *F. asiaticum* (NRRL 6101) genomes (Tralamazza et al., 2019). Genes were predicted with Augustus v.2.5.5 (Stanke & Morgenstern, 2005) based on the pretrained gene prediction database for the *F. graminearum* s.s. genome. Functional gene prediction was performed with Predector v.1.2.7 tool (Jones et al., 2021). Biosynthetic gene clusters (BGC) were predicted with antiSMASH v.7.0 (Blin et al., 2023) and a BGC pangenome was constructed based on the mapping of the biosynthetic-core gene (backbone) against all other genomes. BLASTp v.2.8 (Camacho et al., 2009) local alignment search (Blastp with default parameters) was performed and matches with the highest bitscores were retrieved.

### FGSC phylogenomic analyses

We used single-copy orthologs conserved among all strains to build a phylogenomic tree. Orthology analyses across species were performed with Orthofinder v.2.2 (Emms & Kelly, 2019). Orthologues sequence alignment was performed using MAFFT v.7.3 (Katoh et al., 2017) with parameters –maxiterate 1000 –localpair. We used RAxML v.8.2.12 (Stamatakis, 2014) to construct a maximum-likelihood phylogenetic tree for each alignment with parameters -m PROTGAMMAAUTO and bootstrap of 100 replicates). The final phylogenomic tree was constructed using Astral III v.5.1.1 (C. Zhang et al., 2017) which uses the coalescent model and estimates a species tree given a set of unrooted gene trees. Figtree v.1.4.0 was used for tree visualization (Rambaut A., 2009).

### Seedling infection assay

RNA-seq and metabolome analyses were performed on fungal mycelium grown *in vitro* and *in planta* for five species of the FGSC (*F. graminearum* s.s.*, F. meridionale, F. cortaderiae, F. asiaticum* and *F. austroamericanum*). Each species was grown in a Petri dish containing V8 agar rich medium (10g agar, 30 mM CaCO_3_, 20%, v/v, vegetable juice; Campbell Food, Puurs, Belgium) for four days at 25°C. After culturing, the mycelium was transferred to a 100 ml mung bean liquid mediumClick or tap here to enter text. and agitated at 170 rpm for five days at 25°C. Next, the medium was filtered, and spores counted using a hemocytometer. A 10^6^ per ml spore solution was prepared for the following infection procedure. For RNA-seq in culture strains were inoculated from spore solutions in yeast sucrose agar medium for 72h at 25°C until RNA extraction. For *in planta* assays, wheat coleoptiles with intermediate susceptibility to FHB (cultivar CH Combin, harvest 2018-2019) were infected with each strain individually according to (Zhang et al. 2012). Briefly, wheat seeds were soaked in sterilized water for germination in a culture chamber at 25°C with a 12h white light cycle and 93% humidity. After 3 days of germination, the tip of the coleoptile was cut and 10µl of the spore solution was used for inoculation. Coleoptiles were collected 96h after inoculation (approximately 0.4 mm lesion size) and processed for RNA and metabolome extraction. All *in planta* assays were performed in triplicates and each replicate was composed of a pool of 24 infected seedlings to obtain sufficient material and homogenized infection conditions.

### RNA extraction and sequencing

RNA analyses were performed previously (Tralamazza et al. 2022). Briefly, RNA extraction was performed using the NucleoSpin RNA Plant and Fungi kit (Macherey-Nagel GmbH & Co. KG, Düren, Germany) according to the manufacture’s recommendation. The RNA quality was assessed using a Qubit (ThermoFisher Scientific, Waltham, USA) and an Agilent Bioanalyzer 2100 (Agilent Technologies). The NEB Next® Ultra™ RNA Library Prep kit based on the polyA method was used for RNA library preparation. Samples were sequenced on a NovaSeq 6000 system (Illumina Inc.) and 150 bp paired-end reads were generated. Library preparation and sequencing was performed by Novogene Inc. Illumina raw reads were trimmed and filtered for adapter contamination using Trimmomatic v. 0.32 (parameters: LEADING:3 TRAILING:3 SLIDINGWINDOW:4:15 MINLEN:36) (Bolger et al., 2014). Filtered reads were aligned using Hisat2 v. 2.0.4 with default parameters (Kim et al., 2019) to the *Fusarium graminearum* PH-1 reference genome (King et al., 2017). Mapped transcripts were quantified using HTSeq-count (Anders et al., 2015). Next, reads counts were processed as counts per million (CPM) and normalized by the TMM method using the calcNormFactors function in the R package edgeR (Robinson et al., 2010). To account for gene length, we calculated reads per kilobase per million mapped reads (RPKM) values. Differential gene expression (DGEs) was assessed based on the decideTests function with the default options of the limma v. 3.42.0 R package (Ritchie et al., 2015). DGE was defined as transcripts with a fold-change in expression level greater than 2.0 RPKM and a *q*-value < 0.05. To compare DGEs among FGSC for effects on host infection, we used the curated PHI-base and selected *F. graminearum* s.s. as the focal species. Duplicated gene targets were removed. Pathogenicity-affected phenotypes included increased/reduced virulence, loss of pathogenicity, and lethality.

### Metabolome extraction

We performed an untargeted metabolome analysis of the five FGSC species under *in vitro* culture and *in planta* condition. For *in vitro* conditions, isolates were grown on YSA medium plates (Sigma Aldrich) in triplicates for 4 days at 25°C. For *in planta* analyses, we performed the seedling coleoptile infection assay as previously described with 3-6 biological replicates per isolate. Extractions were performed with 15 mg of material (mycelia or plant coleoptile) in 2 ml screw tubes with two metal beads (3.2mm) each and kept at -80°C. The material was ground in a Qiagen Tissue Lyser at 30Hz for 10 sec. Then, 0.15 ml solution of H_2_O:methanol:formic acid (20:80:0.1, v/v) was added and shaken for 3 min at 30Hz. Tubes were centrifuged at 12’000 g for 2.5 min and the supernatant was collected. Centrifugation was repeated and 100 µL of the supernatant was placed in an HPLC vial for metabolomic analysis.

### Untargeted metabolite profiling

Metabolome analyses were carried out by UHPLC-HRMS using an Acquity UPLC coupled to a Synapt G2 QTOF mass spectrometer (Waters). An Acquity UPLC HSS T3 column (100×2.1mm, 1.8 μm; Waters) was employed at a flow rate of 500 μl/min and maintained at a temperature of 40°C. The following gradient with 0.05% formic acid in water as mobile phase A and 0.05% formic acid in acetonitrile as mobile phase B was applied: 0-100 % B for 10 min, holding at 100% B for 2.0 min, re-equilibration at 0% B for 3.0 min. The injection volume was 2.5 μl. The QTOF was operated in electrospray positive mode using data-independent acquisition (DIA) alternating between two acquisitions functions, one at low and another at high fragmentation energies. Mass spectrometry parameters were as follows: mass range 50-1200 Da, scan time 0.2 s, source temperature 120°C, capillary voltage 2.5 kV, cone voltage 25V, desolvation gas flow and temperature 900 L/h and 400°C, respectively, cone gas flow 20 L/h, collision energy 4 eV (low energy acquisition function) or 15-50 eV (high energy acquisition function). A 500 ng/ml solution of the synthetic peptide leucine-enkephaline was infused constantly into the mass spectrometer as internal reference to ensure accurate mass measurements (<2ppm). Standards for deoxynivalenol, zearalenone and nivalenol were used as reference (Sigma Aldrich, Inc.). Data was recorded by Masslynx v.4.1 (Waters, Inc.). Marker detection was performed using Markerlynx XS (Waters, Inc.) with the following parameters: initial and final retention time 1.5 and 10.0 min, mass range 85-1200 Da, mass window 0.02 Da, retention time window 0.08 min, intensity threshold 500 counts, automatic peak width and peak-to-peak baseline noise calculation, deisotoping applied. Data was mean-centered and Pareto scaled before applying multivariate analyses. Markers of interest were characterized using the mass spectral database spectra database MassBank (https://massbank.eu) and PubChem database.

### Biosynthetic gene cluster deletion assay

*F. graminearum* and *F. cortaderiae* were grown for 4 days in 20 mL in potato-dextrose broth. Genomic DNA was extracted from 100 mg lyophilized tissue applying the FastDNA SPIN Kit for Soil (MP Biomedicals) with the Lysing Matrix E and a Bead Mill 24 Homogenizer (Fisher scientific). For the targeted genes, flanking sequences of between 810-1463 nucleotides were PCR-amplified by the Phusion Hot Start II DNA Polymerase (ThermoFisher, Inc.) following the manufacturer’s guidelines using primers comprising additional 20-nucleotide tails for plasmid assembly. Primers for vector construction Eurofins Genomics (Ebersberg, Germany) were designed using Primer3Plus (Supplementary Table S2). For plasmid construction, primers F071 and F072 were used for amplification of the *hph* gene from pRF-HU2 (Frandsen et al. 2012) comprising the hygromycin resistance gene (*hph*) under control of the *Aspergillus nidulans trpC* promoter and terminator. Primers F073 and F074 were used to amplify the plasmid backbone from pSHUT3-32 (M. R. Nielsen et al., 2019) comprising replication origins for bacteria (oriV, trfA) and yeast (CEN/ARS) together with kanamycin resistance gene (kan) for selection in bacteria and the auxotrophic selection marker URA3 for selection in yeast. Up and Down fragments were evaluated by 1% gel electrophoresis and purified using the QIAquick PCR purification Kit (Qiagen). The *hph* cassette and vector backbone PCR reactions were treated with *Dpn*I (ThermoFisher ER1701) following the manufacturer’s instructions. For vector assembly, 0.2 pmol of each of the four fragments (backbone, *hph* cassette, Up, Down) were mixed for every construct and transformed into competent spheroplasts of *S. cerevisiae* BY4743 using the LiAc/ssDNA/PEG method (Gietz & Schiestl 2007). Overhangs included in primers used for PCR amplification of Up and Down fragments comprised nucleotides identical to the 5’- and 3’-ends of both plasmid backbone and *hph* fragments and acted as template for *in vivo* homologous recombination. The transformation mix was selected for 2 days at 30 ᴼC on Yeast Synthethic Drop-out agar without uracil (Sigma Aldrich Y1501) with added yeast nitrogen base without amino acids (Sigma Aldrich Y0626) and 2 % glucose. From each transformation reaction plate single colonies were inoculated in 5 mL YEPD medium (1 % Yeast extract, 2 % peptone, 2 % glucose) and grown overnight at 30 ᴼC, 180 rpm. The yeast cells were centrifuged, and the pellets resuspended in 250 μl P1 buffer (50 mM Tris-HCl, 10 mM EDTA, 100 μg/mL RNase A), added 5 μl Zymolase solution (10 mg/mL Zymolase 20T from *Arthrobacter luteus* MP Biomedicals, 25 % glycerol, 50 mM Tris-HCl) and incubated at 37 ᴼC for one hour before purifying the plasmid using the QIAprep Spin Miniprep Kit, following the manufacturer’s guidelines. All plasmids were transformed into chemically competent *E. coli* XL-1 Blue cells by heat shock transformation and selected on LB agar containing 50 μg/mL kanamycin. Single colony isolates were streaked on fresh selective plates and the isolated constructs were validated using PCR with primers G056 / G057 to test for presence of the hph gene, and C090/G058 and G059/C093 to confirm the intended recombination between the four fragments (Supplementary Figure S1A).

*Agrobacterium tumefaciens*-mediated transformation (ATMT) was performed as described previously (Malz et al., 2005) with modifications according to (Frandsen et al., 2012). *Agrobacterium* strains carrying pKO-X vectors were inoculated in 10 mL IMAS medium (40 mM MES, 0.2mM acetosyringone, 0.2% glucose, 0.5% glycerol, 11 mM KH_2_PO_4_, 12 mM K_2_HPO_4_, 2.6 mM NaCl, 2 mM MgSO_4_·7H_2_O, 0.44 mM CaCl_2_·2H_2_O, 0.01 mM FeSO_4_·7H_2_O, 3.8 mM (NH_4_)2SO_4_) from overnight cultures in selective LB and grown at 28 ᴼC while shaking at 100 rpm until an OD600 of 0.6-0.7 was reached. 1 mL of IMAS culture was diluted 1:1 with fungal macroconidia resulting in a final concentration of 10^6^ spores /mL. This co-cultivation mix was spread on top black filter papers placed on top of 10 IMAS agar plates, resulting in 2*10^5^ spores per plate. The co-cultivation was incubated at 25 ᴼC in darkness for 3 days before the filters were transferred carefully to fresh Defined Fusarium Medium agar containing 300 μg/mL cefoxitin and 150 μg/mL hygromycin B using sterile tweezers. Filters were transferred to Czapek Dox agar containing 150 μg/mL hygromycin B. All putative transformants were isolated on individual plates and re-streaked twice on selective agar before performing colony PCR evaluating the integrity of the T-DNA inserts (Supplementary Figure S1B).

### Mutant phenotyping assays

Isolates were pre-grown on V8 culture plates for 3 days at 30°C. For *in vitro* mycelia radial growth assays, 5 mm agar plugs were inoculated in rich media (Fusarium complete medium (FCM, (Pontecorvo et al. 1953), V8 and in Minimal medium (Synthetic ICI supplemented with 6 mM glutamine (Geissman et al., 1966) in duplicate. For *in vitro* spore radial growth assays, 5 mm agar plugs were inoculated in 50 mL mung bean broth in the case of Fc and Fg was cultivated in liquid carboxymethyl cellulose (CMC) (Cappellini & Peterson, 1965) in 300 mL Erlenmeyer flasks with baffles were incubated at 160 rpm, at 30 °C for four days. A total of a thousand conidia were inoculated in V8, FCM and ICI media for 4 days at 30°C in the dark. For conidiospore counting, we inoculated two 5 mm agar plugs in 50 mL mung bean broth or CMC for Fc and Fg, respectively in 300 mL Erlenmeyer flasks with baffles at 160 rpm shaking at 30 °C for four days.

### *Fusarium* Head Blight assay

For the infection assay we used dwarf hard red spring wheat USU Apogee wheat (*Triticum aestivum* cv. USU-Apogee; Reg.no CV-840, PI592742) susceptible to FHB. Infection assays were performed as previously described (Studt et al., 2017). Briefly, during the anthesis phase two spikelets of at least five individual wheat heads were inoculated with 1,000 conidia each. As a control, at least three individual wheat ears were treated with sterile water only. After inoculation the treated plantlets were covered with a moistened plastic bag for 24 h to retain high humidity and incubated for 14 days at 20°C, 70 % humidity for 18 h light and 6 h darkness. Infection progress was monitored from third day up to 14 days and FHB symptoms on wheat head were counted.

## RESULTS

### Comparative genomics of the FGSC and gene cluster diversity

We performed integrative analyses to identify major regulatory elements involved in the infection process of species within the FGSC (Supplementary Figure S1C). To establish baseline phylogenetic relationships within the FGSC, we performed phylogenomic reconstruction and comparative genomic analyses based on 16 representatives FGSC genomes and three close members within the *F. sambucinum* species complex (FSAMSC). Genome assemblies ranged in size from 35.8-41.9 Mb (Supplementary Table S1). Orthologues analyses based on BUSCO showed completeness above 98.6% for all assemblies (Supplementary Table S1). We performed phylogenomic reconstruction of the species complex based on 7278 single-copy orthologues (Supplementary Table S3) and conserved within FSAMSC species based on a coalescent model (Figure 1A). Members of the FSAMSC *F. sambucinum, F. culmorum and F. pseudograminearum* were used as outgroups. Members of the FGSC grouped together as a monophyletic clade (Figure 1A). Within the FGSC, tree topologies revealed species falling into three clades consistent with previous multi-locus analyses (Yli-Mattila, 2009). Gene count varied between 11,484-14,145 genes (Figure 1A, Supplementary Table S1), with *F. graminearum* s.s. displaying the highest gene set, at least in part due to the highly complete genome assembly. We found that 71.1% of all orthologues were conserved among FGSC strains (*i.e.* core) and 28.9% were shared among fewer species (*i.e.* accessory) (Figure 1B, Supplementary Table S3).

**Figure 1.**
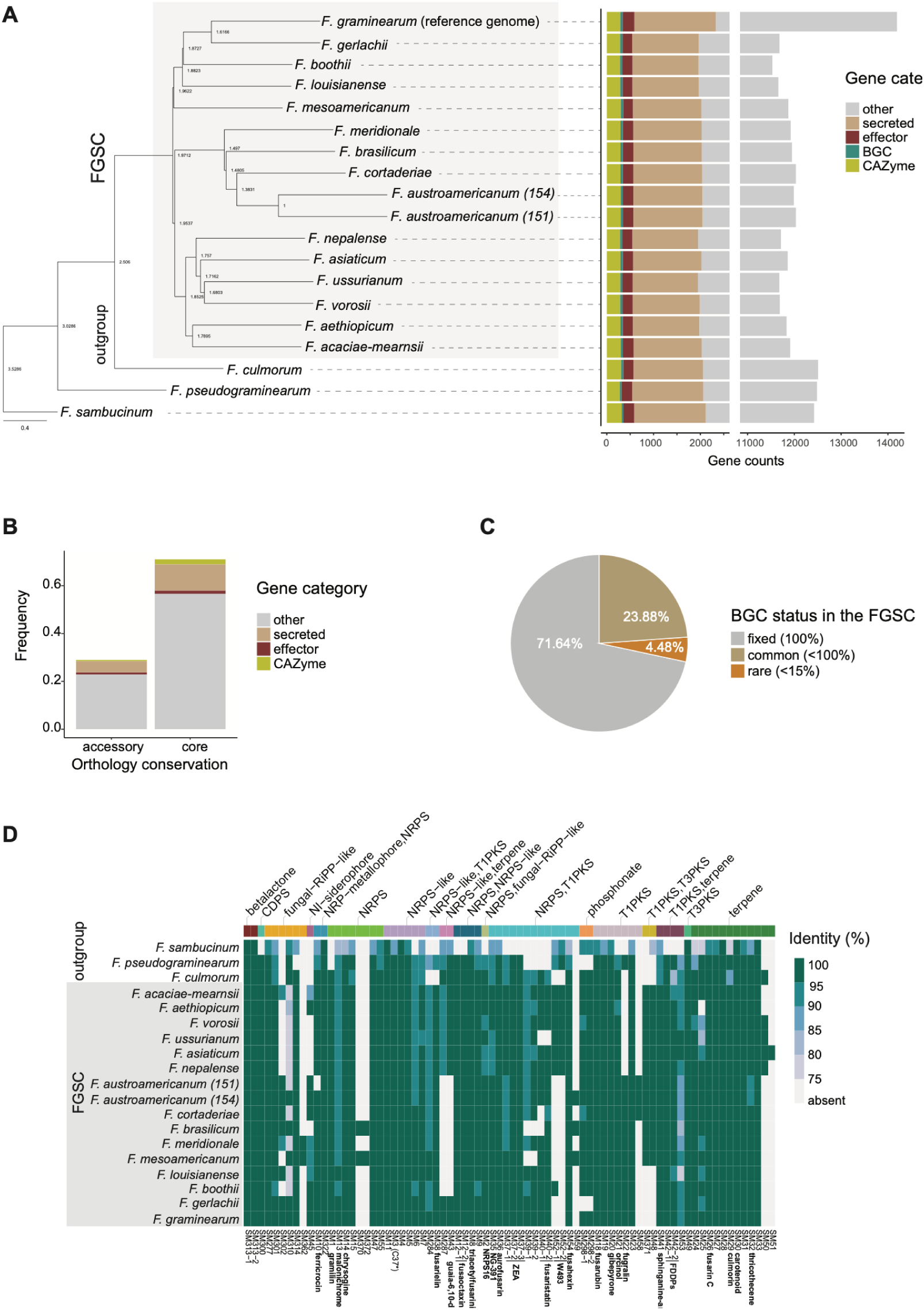
A) Phylogenomic tree of the *Fusarium graminearum* species complex (FGSC). *F. sambucinum, F. culmorum* and *F. pseudograminearum* were used as outgroups. The tree was inferred from a coalescence-based analysis of 7278 single-copy orthologues conserved within thh *Fusarium sambucinum* species complex. Bars correspond to the total gene number. B) Frequency of core and accessory genes within FGSC. Colors correspond to the gene classification based on the Predector tool. C) Biosynthetic gene cluster (BGC) frequency based on conservation shared across FGSC members. D) Heatmap of all predicted BGCs within FGSC. Colors refer to amino acid identity percentage against *F. graminearum* s.s. Top colored bars identify the BGC class. Gray boxes on the right identify members of the FGSC. * refers to Sieber et al (2014) BGC nomenclature.

Gene function analyses supported a conserved virulence toolbox within FGSC species with 60.4%, 69.9% and 82.1% of effector, secretion and CAZyme gene functions, respectively, shared among all species (Figure 1B, Supplementary Table S4). We exhaustively searched for BGCs across all species and found 67 predicted BGCs within the FGSC (Supplementary Table S5). Interestingly, the majority of BGCs (71.64%) are shared among all FGSC species (fixed category, Figure 1C), while 4.48% are found in less than 15% of isolates (rare category, Figure 1C). We found 14.9% of BGCs to be absent in the *F. graminearum* s.s reference genome. The FGSC encoded a functionally diverse set of BGCs, with NRPS-PKS hybrid (n = 13) and terpenes (n = 12) among the most frequent BGC classes identified (Figure 1D). We also found six fungal RIPP-like BGCs within the group. RIPP are a recently discovered class of BGCs, with a potential to reveal exploitable drugs (Vogt & Künzler, 2019), yet their functions within the FGSC remain elusive. The high diversity of BGCs within the FGSC and their conserved nature suggest that the SM machinery may play a conserved role in the host infection process on the small cereal hosts.

### FGSC regulatory strategies during host infection

We performed transcriptomic analyses during wheat coleoptile infection (Figure 2A) and on a nutrient-rich growth medium of five FGSC species (*F. graminearum, F. meridionale, F. cortaderiae, F. austroamericanum* and *F. asiaticum*) that reflect the phylogenetic breadth of the FGSC (Supplementary Table S6). For each sample, 5–55M RNA-seq clean reads were obtained and aligned to the reference genome of *F. graminearum s.s*. Mapped reads varied between 5–23 million among replicates and conditions. A transcriptome-wide principal component analysis (PCA) showed clear separation among FGSC members (Figure 2B, Supplementary Figure 2A). *In planta* replicates of *F. austroamericanum*, *F. meridionale* and *F. cortaderiae* clustered more closely together than the other species (Figure 2B), likely reflecting the phylogenetic signal and possibly adaptation to their shared habitat (*i.e.*, Southern Brazilian wheat crops). *F. asiaticum* showed the most divergent profile, which could be linked to the original host adaptation (*i.e.* maize) compared to the other strains with an original wheat host.

**Figure 2.**
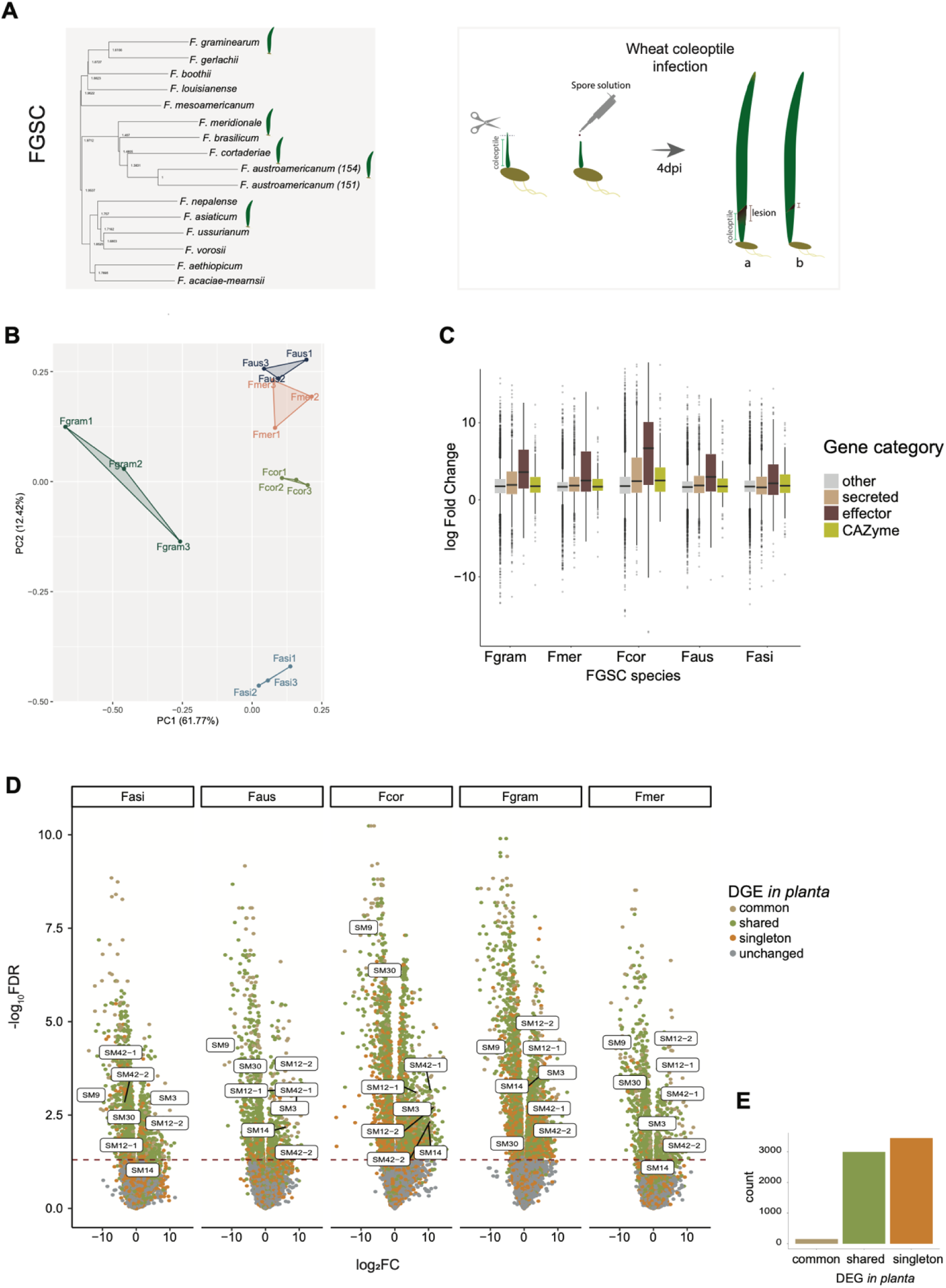
A) FGSC transcriptomic analyses *in planta* and *in vitro*. The leaf symbol indicates species used for the analysis. Left box outline the coleoptile infection procedure used for RNASeq and metabolomic analyses. Dpi refers to days after infection. B) Transcriptomic profiles of FGSC species during host infection based on principal component (PC) analysis. Colors refer to the species (Fgram: *F. graminearum s.s,* Fmer: *F. meridionale,* Fcor: *F. cortaderiae,* Faus: *F. austroamericanum* and Fasi: *F. asiaticum*). Numbers refer to the number of replicates. C) Gene expression fold-change differences between *in planta* and *in vitro* conditions. Values above 1 indicate higher transcription during host infection. Colors refer to gene types. D) Differential gene expression (DGE) between *in planta* and *in vitro* conditions. Red dotted line outline FDR values < 0.05. Colors indicate DGE genes shared between FGSC species. The common category refers to DGEs in all species, shared refers to DGEs shared in less than five species except singletons. Unchanged are genes not differentially expressed between conditions. E) Gene count of each DGE category.

We found that the mean expression of core genes (*in planta* mean = 102.0 RPKM, *in vitro* mean = 99.5 RPKM) was higher compared to accessory genes (*in planta* mean = 41.0 RPKM, *in vitro* mean = 48.2 RPKM; Wilcoxon test, p-value <0.00001; Supplementary Figure 2B). All analyzed FGSC members exhibited upregulation of predicted effectors during infection with *F. cortaderiae* displaying the largest variation (fold change = 4.45) among the group members (Figure 2C). We performed differential gene expression (DGE) analyses to identify infection-upregulated genes compared to the culture condition. To robustly analyze joint gene regulation patterns, we focused on single-copy orthologues conserved across the FGSC (*n* = 7278). We first investigated the degree of shared regulation across FGSC members. We found that 2,993 and 3,446 genes were differentially regulated during infection among some species (*i.e.* shared category) or a single species only (*i.e.* singleton category), respectively (Figure 2D-E). The majority of genes showed no significant infection upregulation among species (mean share = 78.8%; FDR > 0.05). Interestingly, a small proportion of genes (*n* = 155) were differentially regulated *in planta* across all species. This subset of genes showed also among the strongest significant infection upregulation among species (mean = 3.2 -log10(FDR)) compared to other gene categories (Figure 2D; Supplementary Figure 3A). Functional prediction showed that infection-upregulated genes shared among species were particularly enriched for metabolic functions (Supplementary Figure 3B, Supplementary Table S7). We also searched the pathogen–host interaction (PHI)-base database for mutant line reports of *F. graminearum* genes. We found 429 differentially expressed genes of FGSC members with database reports for pathogenicity association (Supplementary Table S8). Most of the shared upregulated genes or singleton upregulated genes were not associated with a PHI-base entry linked to pathogenicity. In contrast, jointly upregulated genes shared among all analyzed FGSC members were 1.8 times more likely to be annotated with a PHI-base pathogenicity association than not (Supplementary Figure S3C). Taken together, our results show that FGSC members share a small core set of shared and jointly upregulated genes with likely pathogenicity contributions.

### Shared metabolic processes during wheat infection within the FGSC

We investigated the BGC expression profiles of all analyzed FGSC species during infection in contrast to culture conditions. We found that the majority of BGCs were actively transcribed (82.1%) in at least one condition and/or species (Supplementary Figure 4A). Clustering analyses showed four dominant BGC gene expression profiles within the FGSC (Figure 3A). A small portion of BGCs (4.9%) were upregulated during *in vitro* conditions compared to *in planta* (*i.e.* profile I; Figure 3A). Approximately 23% of BGCs were highly expressed in both conditions (*i.e.* profile II; Figure 3A), which included the siderophores ferricrocin and triacetylfusarinin. The majority of BGCs (55.7%) show high interspecies variation (profile IV), suggesting species-specific regulation and adaptation.

**Figure 3.**
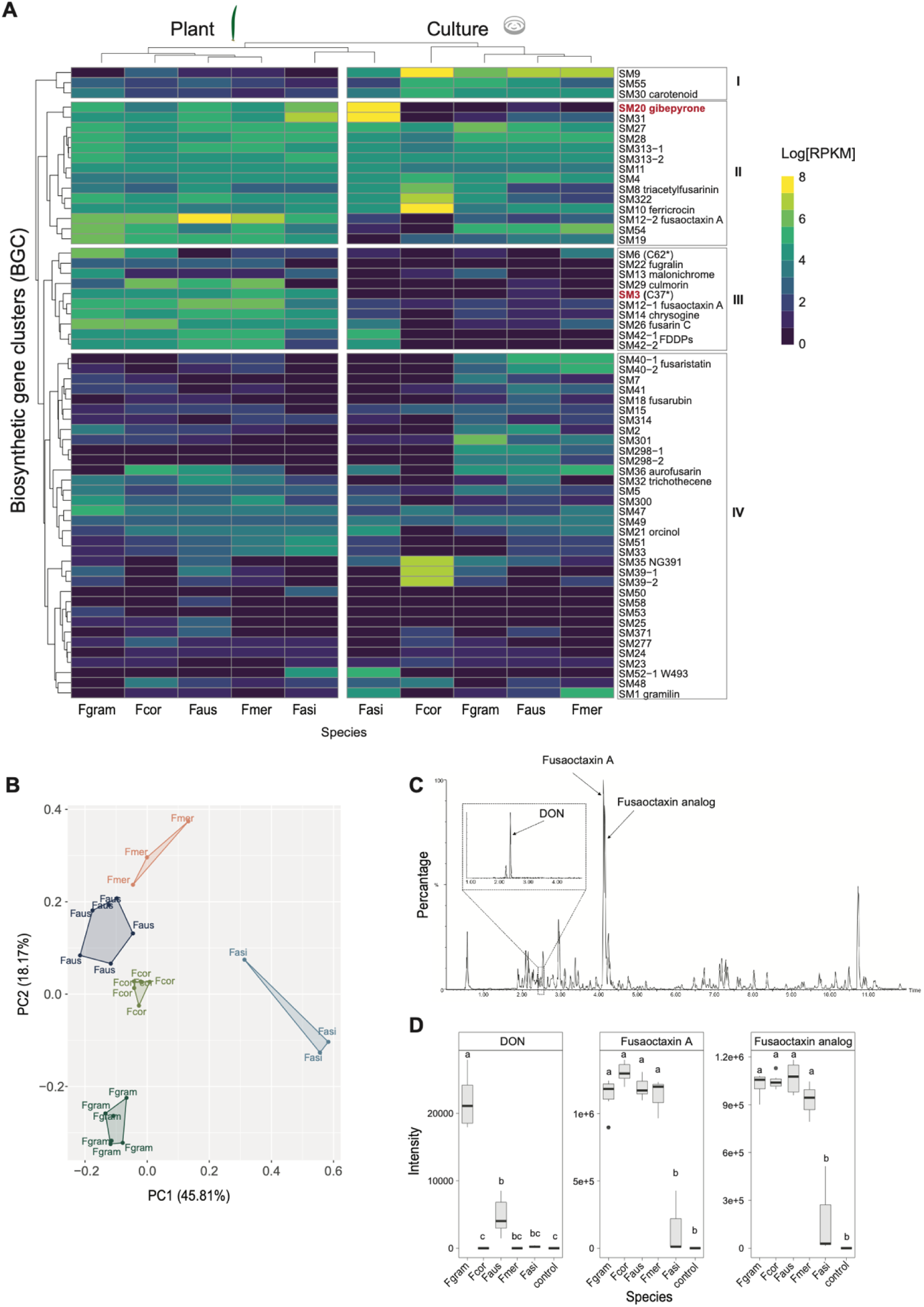
A) BGC expression profiles among FGSC species were analyzed during *in planta* and *in vitro* conditions. The heatmap color represents the RPKM values. The numbers I, II, III, and IV to the right indicate the hierarchical clustering of BGC expression profiles. * refers to Sieber et al (2014) BGC nomenclature. B) Metabolomic profiles of FGSC species during host infection were analyzed based on principal component (PC) analysis. Colors refer to the species (Fgram: *F. graminearum s.s,* Fmer: *F. meridionale,* Fcor: *F. cortaderiae,* Faus: *F. austroamericanum* and Fasi: *F. asiaticum*). C) Metabolomic chromatogram of a representative sample (*F. graminearum* s.s during plant infection) highlighting the major virulence factors deoxynivalenol, fusaoctaxin A and analog. D) Metabolite intensity of deoxynivalenol, Fusaoctaxin A and analog across FGSC species during host infection. Different letters above the boxplot identify significantly different groups according to an ANOVA and Tukey test.

We focused on highly active BGCs during the infection process across all species (*i.e.* profile III; Figure 3A), as the produced SMs could play a role as virulence factors. We identified nine BGCs (16.5%) matching this profile, including frugalin, the mycotoxin fusarin C, culmorin and the known virulence factor fusaoctaxin A. SM42 (PKS15) is known to produce fungal decalin-containing diterpenoid pyrones (FDDPs) (C. Hicks et al., 2023) and showed one of the strongest DGE signals during wheat infection. Among profile III, we found SM3 (C37; Sieber et al 2014) and SM6 (C62), two non-canonical NRPS-like BGCs. To potentially match BGC upregulation with candidate SMs, we performed untargeted metabolomic analyses under the same conditions as the transcriptome analyses (Supplementary Table S9). We found that the overall metabolomic profiles of FGSC species matched their expression profile leading to clustering of phylogenetic closer species (*i.e. F. cortaderiae, F. austroamericanum* and *F. meridionale*; Figure 3B, Supplementary Figure S4B). Hence, transcriptional regulation and SM production both likely share a common phylogenetic signal. Next, we focused on BGCs and associated virulence factors. We found that SM37 deoxynivalenol (DON) was most abundantly produced by *F. graminearum* s.s. followed by *F. austroamericanum*. No production was detected in the other FGSC members (Figure 3D). Fusaoctaxin A (NRPS5/NRPS9) showed the highest metabolite peak intensities across samples together with *F. asiaticum* (Figure 3C). The strong intensity of fusaoctaxin A is likely explained by the condition used (*i.e.* coleoptile infection process), since the metabolite is required for proper cell-to-cell invasion of wheat by *F. graminearum*. In contrast, DON is required for successful mature wheat spikelet infection (Bai, Desjardins, and Plattner 2002). Our data suggest that fusaoctaxin A is likely a key metabolite not only for *F. graminearum* s.s. but several additional FGSC members during host infection. Based on the same infection-upregulation profiling (*i.e.* profile III; Figure 3A) and DGE signal (Figure 2D), we identified SM42 (PKS15), the uncharacterized SM3 (C37) and SM6 (C62) as strong candidates for contributing to pathogenicity.

### Gibepyrone impact on host infection

Gibepyrone (SM20 - PKS8) marginally falling into profile II shows a largely consistent upregulation during infection except for *F. asiaticum* where culture medium induces the highest transcription (Figure 3A). To test the potential role of the metabolite, we deleted the core gene *PKS8* of the gibepyrone cluster in *F. cortaderiae* (FcWT) (Supplementary Figure S1AB). We found no apparent growth (Supplementary Figure S5) or macromorphological anomalies between mutants lacking *PKS8* (Δ*PKS8FC*T1 and *ΔPKS8FCT2*) compared to the wild-type strain (FcWT) in the three tested culture media (*i.e.* FCM, ICI and V8). Next, we examined the impact of gibepyrone production defects during wheat infection. Based on the coleoptile infection method, we found that the mutant *ΔPKS8FCT1* caused a larger lesion area (mean = 5.5 mm, *p* < 0.05) than the wild-type strain (mean = 5.0 mm; Figure 4A). However, no significant difference between mutants and wild-type was found based on the wheat spikelet method, which mimics FHB disease (Figure 4B). Spore production counts showed that the mutant *ΔPKS8FCT2* produced more spores (mean = 3×10^7^ spores, *p*-value <0.05) compared to the wild type (mean = 2×10^7^ spores). Taken together, our results indicate that gibepyrone has no direct impact on cell-to-cell or spikelet wheat infection in *F. cortaderiae*. The larger lesion symptoms and increased spore production displayed by the inactive BGC may be a reflection of the avoided costs of producing the metabolite.

**Figure 4.**
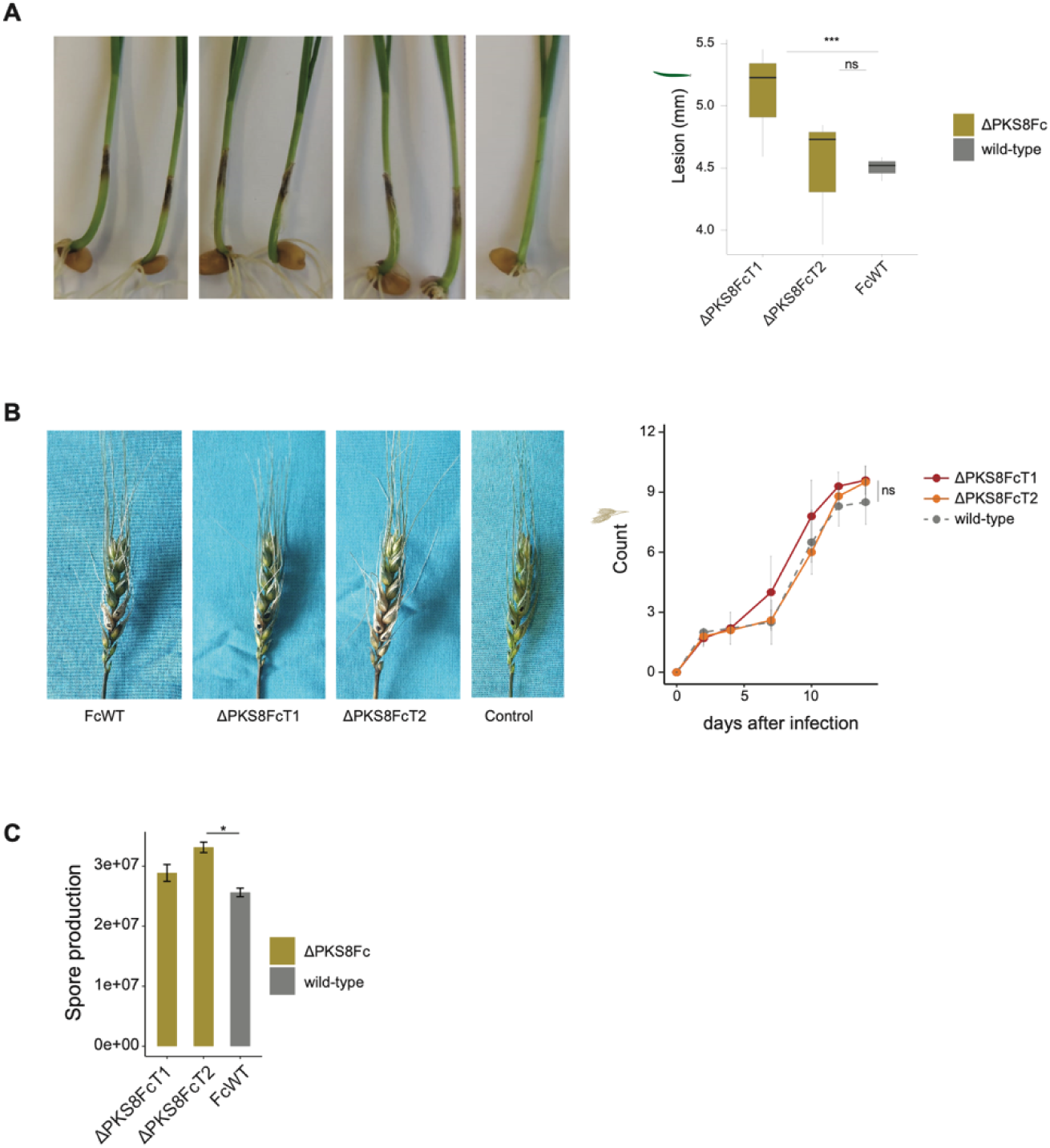
Evluation of gibepyrone effects in *F. cortaderiae.* A) Representative images of wheat seedlings at 4 days after coleoptile inoculation with *F. cortaderiae* wild-type strain (FcWT) and PKS8 core gene deletion mutants (ΔPKS8Fc). The control was inoculated with water. The right boxplot refers to the distribution of lesion sizes produced by the different genotypes. B) Representative images of wheat spikes at 14 days after inoculation with *F. cortaderiae* wild-type strain (FcWT) and PKS8 core gene deletion mutants (ΔPKS8Fc). The control was inoculated with water. The right plot refers to the number of infected wheat heads per condition. C) Number of conidiospores produced by the tested strains. Wilcoxon tests with adjusted *p*-values (Holm method). ns: p > 0.05, *p < 0.05, **p < 0.01, ***p < 0.001, **** p <0.0001.

### An infection role for SM3 in the FGSC

Next, we focused on SM3 BGC shared among all FGSC members (Figure 1B). SM3 is encoded on chromosome 4 with a predicted size of 59.8 kb comprising 23 genes. Our transcriptomic analyses showed that SM3 was one of the top shared and jointly upregulated BGCs during host infection (Figure 2D, 3A). Synteny analyses across FGSC showed that SM3 shares a conserved region among all members comprising five genes, including the core-biosynthetic FGRAMPH1_G22333 (FGSG_06462) and a transporter FGRAMPH1_G22337 gene (Figure 5A, Supplementary Table S10). Flanking regions showed overall high synteny across species, with some segregating structural variation (*i.e.* duplications and translocations). Rearrangements were particularly apparent in *F. mesoamericanum* and *F. acaciae-mearnsii* species (Figure 5A). Mean gene expression among FGSC members during infection showed active transcription mostly in the conserved syntenic region, which could indicate that the functional BGC is smaller than predicted *in silico* (Figure 5B). We investigated SM3 for links to pathogenicity during wheat infection. The non-canonical BGC encodes a NRPS-like biosynthetic core FGRAMPH1_G22333 constitued by a single AMP-binding and PP-binding domain module (Figure 6A). We deleted the core gene FGRAMPH1_G22333 (Figure 6A, Supplementary Figure S1AB) of *F. graminearum s.s.* (mutants *ΔSM3FgT1* and *ΔSM3FgT2*). Morphology of Δ*SM3Fg* mutants showed a denser aerial mycelium (Figure 6B). We assessed radial hyphal growth based on conidia and mycelia under rich culture media (*i.e.* V8 and FCM) and minimal medium (i.e. ICI; Figure 6B). We found that under nutrient starvation (*i.e.* minimal medium) mutants grow faster than the wild-type strain FgWT, while on rich media no difference was observed (Figure 6B). We found that production of conidiospores was strongly hampered in the *ΔSM3Fg* mutants (mean = 1×10^6^ spores; *p*-value < 0.00001; Figure 6C) compared to wild-type (mean = 4×10^7^ spores). We then investigated the impact of SM3 on host infection. We found that during coleoptile infection the mutants significantly lost (mean = 0.5 mm; p-value < 0.0001) the ability to produce lesions compared to the wild strain (mean = 4 mm) (Supplementary Figure 6A). We also observed a significant loss of pathogenicity in the mutants (mean at 14 dpi = 6.4 infected heads) compared to the wild type (mean = 10.1 infected heads) during wheat spikelet infection (Figure 6D). Untargeted metabolomics analyses (Supplementary Table S11) revealed overall distinct metabolic profiles of the mutants compared to the wild-type (Figure 6E, Supplementary Figure 6). We found no significant differences between the major virulence factors deoxynivalenol and fusaoctaxin A under the tested conditions (Figure 6E, Supplementary Figure 6). This indicates that the loss of pathogenicity is independent of previously characterized virulence factors. Taken together, we discovered a shared BGC with a strong impact on pathogenicity on wheat, morphological differentiation and significant decrease of asexual conidiation.

**Figure 5.**
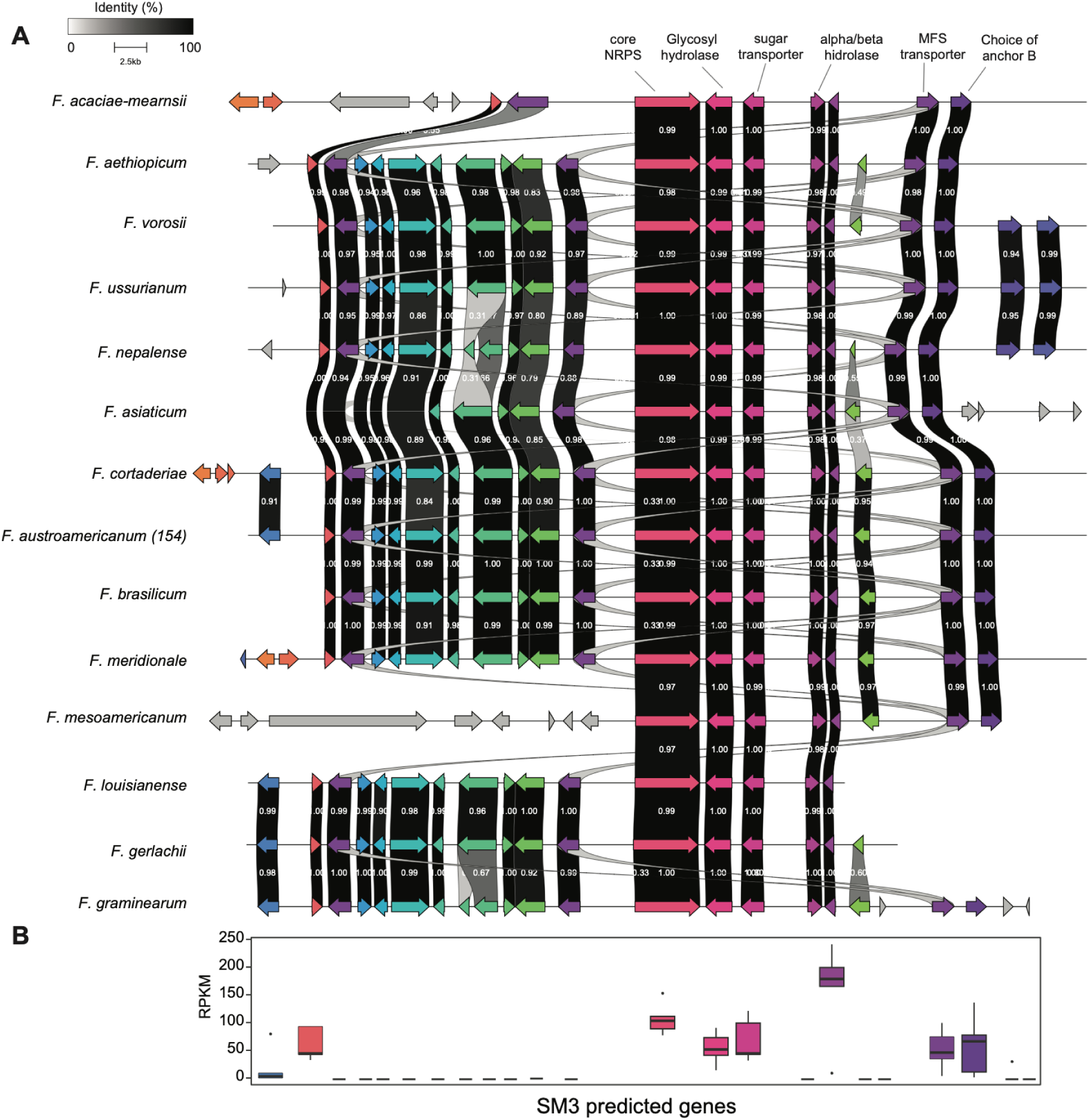
Synteny plot of the BGC SM3 chromosomal region contrasted between FGSC species. Colored arrows identify genes. The gray scale indicates the degree of amino acid identity. B) Mean expression values of the genes encoded by the BGC SM3 among FGSC species during host infection.

**Figure 6.**
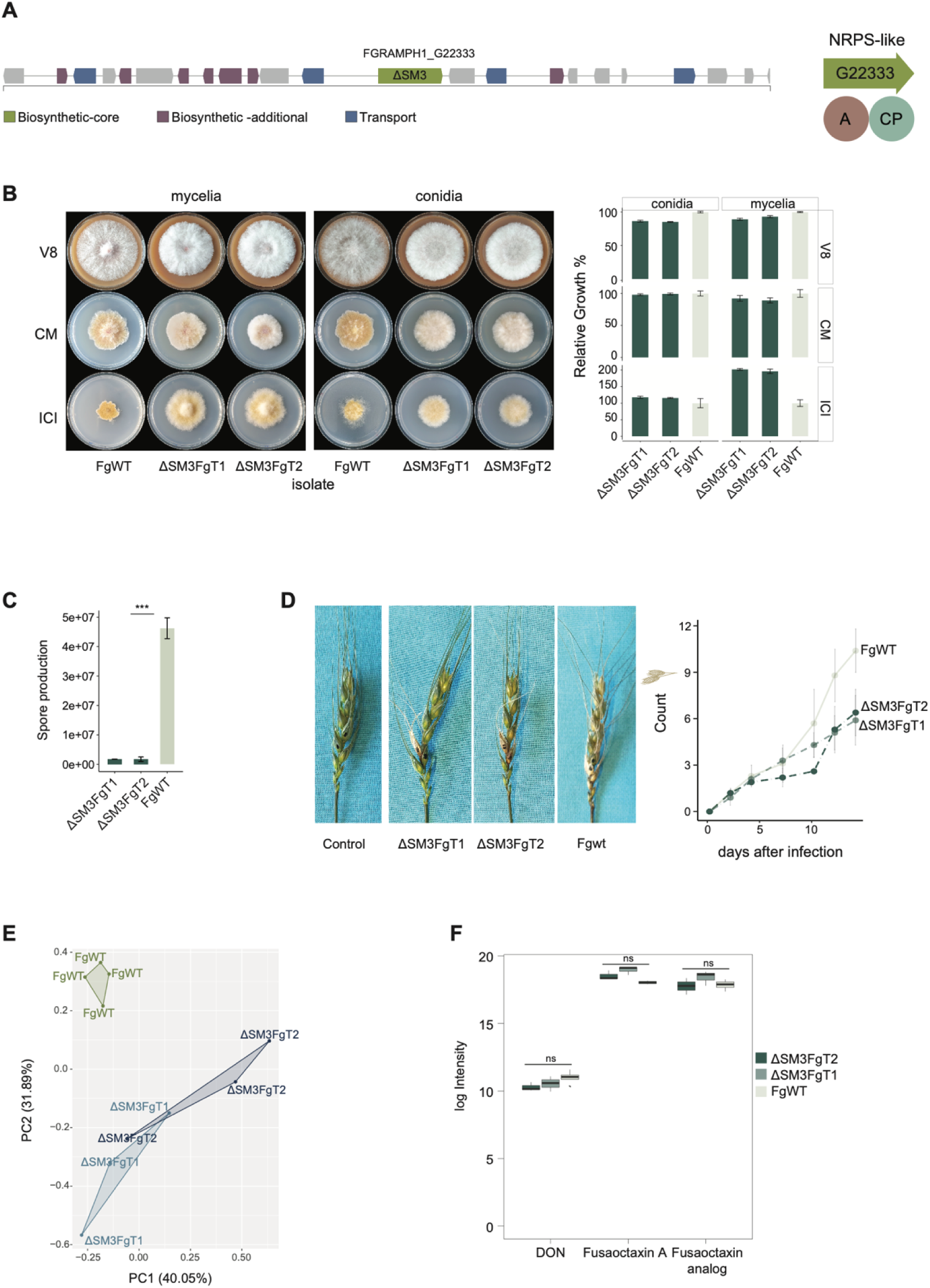
SM3 deletion hampers *F. graminearum* pathogenicity on wheat. A) The biosynthetic-core gene FGRAMPH1_G22333 (SM3) and adjacent genes possibly comprising the SM3 biosynthetic gene cluster based on *in silico* prediction. On the right, the predicted module domains of the biosynthetic NRPS-like gene are shown. “A” refers to AMP-binding and CP to PP-binding domain. B) Representative images of 5-day old cultures of the *F. graminearum* wild-type strain (FgWT) and Δ*SM3Fg* deletion mutants. The right plot refers to the relative fungal growth in each tested culture medium. C) Number of conidiospores produced by the tested strains. D) Representative images of wheat spikes at 14 days after inoculation with *F. graminearum* wild-type strain (FgWT) and PKS8 deletion mutants (ΔSM3Fg). The control was inoculated with water. The right plot refers to the number of infected wheat heads per condition. Wilcoxon tests with adjusted *p*-values (Holm method). Ns: p > 0.05, *p < 0.05, **p < 0.01, ***p < 0.001, **** p <0.0001. E) Metabolomic profile of wild-type (FgWT) and deletion mutant strains (ΔSM3Fg) during host infection based on a principal component (PC) analysis. Colors refer to the different strains. F) Metabolite intensity of deoxynivalenol, fusaoctaxin A and analog in the wild-type (FgWT) and mutant strains (ΔSM3Fg) during host infection. Wilcoxon tests with adjusted *p*-values (Holm method). Ns: p> 0.05.

## DISCUSSION

We performed comparative genomics and transcriptomics in the major fungal pathogen group *Fusarium graminearum* species complex to assess shared and species-specific regulatory dynamics during host infection. We exhaustively identified the regulatory landscape of SMs and shared modulation across species, which allowed us to functionally characterize a new BGC acting as a virulence factor during wheat infection.

Comparative transcriptomics studies often focus on different conditions in a single species (Gay et al., 2021; Kazan & Gardiner, 2018). Here, we analyzed the regulatory landscape of five species of the FGSC during host infection and *in vitro* conditions. We found that closely related species (i.e. *F. austroamericanum, F. cortaderiae* and *F. meridionale*) displayed more similar genome-wide regulation and metabolite production than other species, reflecting the phylogenetic signal within the group (J. C. Nielsen et al., 2019). We show that most conserved orthologues undergo species-specific transcription regulation. The polymorphic regulation among FGSC member is potentially a consequence of adaptation by individual species geographic (van der Lee et al., 2015) and host niches (Amarasinghe et al., 2019; Machado et al., 2021; H. Zhang et al., 2012). In contrast, the small percentage of orthologues jointly upregulated during the infection process (*i.e.* common DGE category) may represent an ancestral trait shared by FGSC members to infect cereal hosts (Blasdel et al., 2017; Miguel-Rojas et al., 2023). The common set of DGE was enriched for metabolic processes and included several BGCs consistent with expectations for many other molecular host-pathogen interactions (Li et al., 2020). Integrating functional studies reported in the PHI-base revealed that jointly upregulated genes more likely play a role in pathogenicity in contrast to genes upregulated in a species-specific manner. Our results indicate that FGSC member retained a conserved toolbox of infection-regulated genes.

Fungal genomes often encode diverse sets of BGCs, yet metabolic products and functions remain largely unknown (Rokas et al., 2020). Species-specific BGCs or silenced biosynthetic pathways can pose significant challenges for functional discovery (Keller, 2015). BGCs encoded by FGSC members were following four distinct expression profiles. We found that upregulation in culture medium compared to host infection presented the rarest BGC expression profile across species. Under laboratory conditions, epigenetic silencing and lack of stimuli are likely factors rendering BGCs transcriptionally silent (Keller, 2019). In contrast, BGCs constitutively expressed across conditions were more abundant and underpin production of gibepyrone and the siderophores ferricrocin and triacetylfusarinin (Oide et al., 2015). Siderophores play critical roles microbial development and pathogenicity under nutrient limitation (Miethke & Marahiel, 2007) and manipulation of siderophores can offer opportunities for pathogen management (Somu et al., 2006). Mutant analyses of gibepyrone BGC deletions revealed no noticeable effects on the infection process. Little is known about the metabolite cellular function apart from moderate antimicrobial (Barrero et al., 1993) and parasitic activity (Bogner et al., 2017). *In vitro* analyses showed that the metabolite is toxic for the producer *F. fujikuroi* (Janevska et al., 2016) yet more detailed investigations of gibepyrone effects are needed.

We also focused on SMs produced by infection-upregulated BGCs. Among these clusters, our strategy identified multiple BGCs associated with known metabolies. Culmorin is a mycotoxin able to repress deoxynivalenol detoxication in plants (Woelflingseder et al., 2019). The BGC underpinning production of fusarin C, which is a metabolite with unknown function during infection is linked though to esophageal and breast cancer progression (Sondergaard et al., 2011). Frugalin (PKS2) is an upregulated polyketide BGC with deletion mutants showing decreased mycelial growth (Gaffoor et al. 2005) and sporulation, with no effect on FHB symptoms (Severinsen et al., 2023). Yet the highly expression of the BGC during coleoptile infection suggests that the SM could be linked to other form of infection. Similarly to the strongly upregulated fusaoctaxin A (NRPS5/NRPS9) virulence factor, essential for host cell-to-cell penetration in *F. graminearum* (Jia et al., 2019). SM42 (PKS15) is linked to FDDP production (C. Hicks et al., 2023), yet the BGC function remains elusive. Our DGE analyses suggest that the PKS15 plays a significant role during the infection process of FGSC. We found that the NRPS-like SM3(C37) and SM6(C62) were among the most conserved upregulation profiles in the FGSC consistent with an important role during infection. Taken together, the comparative infection transcriptome analyses of the FGSC clarified expression profiles of known virulence factors and new, non-canonical metabolic targets for further investigation.

We found that the biosynthetic gene cluster SM3 can act as a virulence factor of *F. graminearum* s.s. during plant infection. The BGC was first identified in *F. graminearum* s.s. (Sieber et al., 2014) and showed high conservation (>90% amino acid identity) within the FGSC and the related FSSC (Tralamazza et al. 2019). The core-biosynthetic gene of the cluster is highly expressed during host infection and co-regulated across multiple species within the FGSC while inactive *in vitro* culture supporting the involvement of SM3 in virulence functions. Metabolic synthesis can be costly (Rokas et al., 2018) and exhibits high variability within and between species at the level of regulatory control (Niehaus et al., 2017). We found that deletion of the core NRPS gene of BGC SM3 significantly impaired the pathogenicity process during wheat coleoptile and FHB infection. The SMs deoxynivalenol and fusaoctaxin A play a key role in virulence of *F. graminearum* on wheat (Jia et al., 2019; Walter et al., 2010). We show that production of these metabolites was not affected by the inactivation of the SM3-key enzyme suggesting regulatory independence between different BGCs conferring virulence (Bergmann et al., 2010). Beyond the impact of pathogenicity, deletion of the SM3 NRPS core gene also hampered asexual spore production in the pathogen. Pleiotropic effects between specialized metabolism and development are likely factors in this and are consistent with observations in other fungi (Calvo et al., 2002). The close relationship between virulence and fungal development after the inactivation of SM3 core gene may be due to a shared regulatory pathway between the two processes (Shimizu & Keller, 2001). This may be similar to how the regulator *flbA* controls the production of the carcinogenic SM sterigmatocystin and asexual sporulation in *A. nidulans* (Tsai et al., 1998). In *Chaetomium globosum* the gene *CgXpp1* acts as a negative regulator of the antitumor chaetoglobosin A metabolite and positive regulator of spore production and cell attenuation (Zhao et al., 2022). Another possibility is that SM3 might stimulate the sporogenic process in FGSC. In *F. graminearum* the estrogenic mycotoxin zearalenone enhances perithecial production during *in vitro* growth (Mirocha & Swanson, 1983; Wolf & Mirocha, 2011) although no direct impact on pathogenicity has been established for this mycotoxin.

Our comparative infection transcriptome and gene cluster analyses of the FGSC represent an important step towards a fuller understanding of SMs mediating cereal infections by *Fusarium* species. The SMs described here may also serve in future toxin screenings. We expect that comparative transcriptomics is a fruitful approach to unravel yet unknown pathogenicity factors in other plant-pathogen systems. Pathosystems with similar or identical hosts colonized by a group of related fungal pathogens are particularly suited given the recent diversification of virulence functions. Dissecting both conserved gene functions as well as joint regulation should be an effective approach to identify core pathogenic toolboxes employed by pathogens.

## Acknowledgements

Anna K. Atanasoff-Kardjalieff was affiliated with the AgriGenomics Doctoral School at the University of Natural Resources and Life Sciences, Vienna (BOKU). We are grateful for comments by Ana Margarida Sampaio on an earlier version of this manuscript.

## Data availability

RNA-Seq raw reads are available from the NCBI SRA database under the accession number PRJNA542165.

## REFERENCES

1. Amarasinghe, C., Sharanowski, B., Fernando, W. G. D., Amarasinghe, C., Sharanowski, B., & Fernando, W. G. D. (2019). Molecular Phylogenetic Relationships, Trichothecene Chemotype Diversity and Aggressiveness of Strains in a Global Collection of Fusarium graminearum Species. Toxins, 11(5), 263. 10.3390/toxins11050263

2. Amos, G. C. A., Awakawa, T., Tuttle, R. N., Letzel, A. C., Kim, M. C., Kudo, Y., Fenical, W., Moore, B. S., & Jensen, P. R. (2017). Comparative transcriptomics as a guide to natural product discovery and biosynthetic gene cluster functionality. Proceedings of the National Academy of Sciences of the United States of America, 114(52), E11121–E11130. 10.1073/PNAS.1714381115/SUPPL_FILE/PNAS.1714381115.SAPP.PDF

3. Anders, S., Pyl, P. T., & Huber, W. (2015). HTSeq--a Python framework to work with high-throughput sequencing data. Bioinformatics, 31(2), 166–169. 10.1093/bioinformatics/btu638

4. Arulprakasam, K. R., & Dharumadurai, D. (2021). Genome mining of biosynthetic gene clusters intended for secondary metabolites conservation in actinobacteria. Microbial Pathogenesis, 161, 105252. 10.1016/J.MICPATH.2021.105252

5. Atanasoff-Kardjalieff, A. K., & Studt, L. (2022). Secondary Metabolite Gene Regulation in Mycotoxigenic Fusarium Species: A Focus on Chromatin. In Toxins (Vol. 14). MDPI. 10.3390/toxins14020096

6. Bai, G. H., Desjardins, A. E., & Plattner, R. D. (2002). Deoxynivalenol-nonproducing Fusarium graminearum causes initial infection, but does not cause disease spread in wheat spikes. Mycopathologia, 153(2), 91–98. 10.1023/A:1014419323550

7. Barrero, A. F., Okra, J. E., Herrador, M. M., Cabrera, E., Sanchez, J. F., Quilez, J. F., Rojas, F. J., & Reyes, J. F. (1993). *Gibepyrones: a-Pyrones from Gibbereh fujikuroi* (Vol. 49, Issue 1).

8. Bergmann, S., Funk, A. N., Scherlach, K., Schroeckh, V., Shelest, E., Horn, U., Hertweck, C., & Brakhage, A. A. (2010). Activation of a silent fungal polyketide biosynthesis pathway through regulatory cross talk with a cryptic nonribosomal peptide synthetase gene cluster. Applied and Environmental Microbiology, 76(24), 8143–8149. 10.1128/AEM.00683-10/SUPPL_FILE/SUPPLEMENTARY_TABLE_2.ZIP

9. Blasdel, B. G., Chevallereau, A., Monot, M., Lavigne, R., & Debarbieux, L. (2017). Comparative transcriptomics analyses reveal the conservation of an ancestral infectious strategy in two bacteriophage genera. The ISME Journal, 11(9), 1988–1996. 10.1038/ISMEJ.2017.63

10. Blin, K., Shaw, S., Augustijn, H. E., Reitz, Z. L., Biermann, F., Alanjary, M., Fetter, A., Terlouw, B. R., Metcalf, W. W., Helfrich, E. J. N., van Wezel, G. P., Medema, M. H., & Weber, T. (2023). antiSMASH 7.0: new and improved predictions for detection, regulation, chemical structures and visualisation. Nucleic Acids Research, 51(W1), W46–W50. 10.1093/NAR/GKAD344

11. Bogner, C. W., Kamdem, R. S. T., Sichtermann, G., Matthäus, C., Hölscher, D., Popp, J., Proksch, P., Grundler, F. M. W., & Schouten, A. (2017). Bioactive secondary metabolites with multiple activities from a fungal endophyte. Microbial Biotechnology, 10, 175–188. 10.1111/1751-7915.12467

12. Bolger, A. M., Lohse, M., & Usadel, B. (2014). Trimmomatic: a flexible trimmer for Illumina sequence data. Bioinformatics, 30(15), 2114–2120. 10.1093/bioinformatics/btu170

13. Boutigny, A. L., Ward, T. J., Van Coller, G. J., Flett, B., Lamprecht, S. C., O’Donnell, K., & Viljoen, A. (2011). Analysis of the Fusarium graminearum species complex from wheat, barley and maize in South Africa provides evidence of species-specific differences in host preference. Fungal Genetics and Biology, 48(9), 914–920. 10.1016/J.FGB.2011.05.005

14. Brauer, E. K., Subramaniam, R., & Harris, L. J. (2020). Regulation and dynamics of gene expression during the life cycle of fusarium graminearum. Phytopathology, 110(8), 1368–1374. 10.1094/PHYTO-03-20-0080-IA/ASSET/IMAGES/LARGE/PHYTO-03-20-0080-IAT1.JPEG

15. Broschla, G., Ransom, R., Lechnerla, T., Walton, J. D., & Loidlai’, P. (1995). Inhibition of maize histone deacetylases by HC toxin, the host-selective toxin of Cochliobolus carbonum. The Plant Cell, 7(11), 1941–1950. 10.1105/TPC.7.11.1941

16. Brown, D. W., Kim, H. S., McGovern, A. E., Probyn, C. E., & Proctor, R. H. (2022). Genus-wide analysis of Fusarium polyketide synthases reveals broad chemical potential. Fungal Genetics and Biology, 160, 103696. 10.1016/J.FGB.2022.103696

17. Brown, D. W., & Proctor, R. H. (2016). Insights into natural products biosynthesis from analysis of 490 polyketide synthases from Fusarium. Fungal Genetics and Biology, 89, 37–51. 10.1016/J.FGB.2016.01.008

18. Bruno, M., Dewi, I. M. W., Matzaraki, V., ter Horst, R., Pekmezovic, M., Rösler, B., Groh, L., Röring, R. J., Kumar, V., Li, Y., Carvalho, A., Netea, M. G., Latgé, J. P., Gresnigt, M. S., & van de Veerdonk, F. L. (2021). Comparative host transcriptome in response to pathogenic fungi identifies common and species-specific transcriptional antifungal host response pathways. Computational and Structural Biotechnology Journal, 19, 647–663. 10.1016/J.CSBJ.2020.12.036

19. Calvo, A. M., Wilson, R. A., Bok, J. W., & Keller, N. P. (2002). Relationship between Secondary Metabolism and Fungal Development. Microbiology and Molecular Biology Reviews, 66(3), 447–459. 10.1128/mmbr.66.3.447-459.2002

20. Camacho, C., Coulouris, G., Avagyan, V., Ma, N., Papadopoulos, J., Bealer, K., & Madden, T. L. (2009). BLAST+: Architecture and applications. BMC Bioinformatics, 10. 10.1186/1471-2105-10-421

21. Cappellini, R. A., & Peterson, J. L. (1965). Macroconidium Formation in Submerged Cultures by a Nonsporulating Strain of Gibberella Zeae. Mycologia, 57, 962–966.

22. Cervini, C., Verheecke-Vaessen, C., Ferrara, M., García-Cela, E., Magistà, D., Medina, A., Gallo, A., Magan, N., & Perrone, G. (2021). Interacting climate change factors (CO2 and temperature cycles) effects on growth, secondary metabolite gene expression and phenotypic ochratoxin A production by Aspergillus carbonarius strains on a grape-based matrix. Fungal Biology, 125(2), 115–122. 10.1016/J.FUNBIO.2019.11.001

23. Cordero, R. J. B., & Casadevall, A. (2017). Functions of fungal melanin beyond virulence. Fungal Biology Reviews, 31(2), 99–112. 10.1016/J.FBR.2016.12.003

24. de Chaves, M. A., Reginatto, P., da Costa, B. S., de Paschoal, R. I., Teixeira, M. L., & Fuentefria, A. M. (2022). Fungicide Resistance in Fusarium graminearum Species Complex. Current Microbiology 2022 79:2, 79(2), 1–9. 10.1007/S00284-021-02759-4

25. De Jonge, R., Ebert, M. K., Huitt-Roehl, C. R., Pal, P., Suttle, J. C., Spanner, R. E., Neubauer, J. D., Jurick, W. M., Stott, K. A., Secor, G. A., Thomma, B. P. H. J., De Peer, Y. Van, Townsend, C. A., & Bolton, M. D. (2018). Gene cluster conservation provides insight into cercosporin biosynthesis and extends production to the genus Colletotrichum. Proceedings of the National Academy of Sciences of the United States of America, 115(24), E5459–E5466. 10.1073/PNAS.1712798115/SUPPL_FILE/PNAS.1712798115.SD01.XLSX

26. Deighton, N., Muckenschnabel, I., Colmenares, A. J., Collado, I. G., & Williamson, B. (2001). Botrydial is produced in plant tissues infected by Botrytis cinerea. Phytochemistry, 57(5), 689– 692. 10.1016/S0031-9422(01)00088-7

27. Drott, M. T., Rush, T. A., Satterlee, T. R., Giannone, R. J., Abraham, P. E., Greco, C., Venkatesh, N., Skerker, J. M., Louise Glass, N., Labbé, J. L., Milgroom, M. G., & Keller, N. P. (2021). Microevolution in the pansecondary metabolome of Aspergillus flavus and its potential macroevolutionary implications for filamentous fungi. Proceedings of the National Academy of Sciences of the United States of America, 118(21), e2021683118. 10.1073/PNAS.2021683118/SUPPL_FILE/PNAS.2021683118.SAPP.PDF

28. Ebert, M. K., Spanner, R. E., de Jonge, R., Smith, D. J., Holthusen, J., Secor, G. A., Thomma, B. P. H. J., & Bolton, M. D. (2019). Gene cluster conservation identifies melanin and perylenequinone biosynthesis pathways in multiple plant pathogenic fungi. Environmental Microbiology, 21(3), 913–927. 10.1111/1462-2920.14475

29. Emms, D. M., & Kelly, S. (2019). OrthoFinder: Phylogenetic orthology inference for comparative genomics. Genome Biology, 20(1), 1–14. 10.1186/s13059-019-1832-y

30. Espeso, E. A., & Peñalva, M. A. (1996). Three binding sites for the Aspergillus nidulans PacC zinc-finger transcription factor are necessary and sufficient for regulation by ambient pH of the isopenicillin N synthase gene promoter. Journal of Biological Chemistry, 271(46), 28825– 28830. 10.1074/jbc.271.46.28825

31. Frandsen, R. J. N., Frandsen, M., & Giese, H. (2012). Targeted gene replacement in fungal pathogens via agrobacterium tumefaciens-mediated transformation. Methods in Molecular Biology, 835, 17–45. 10.1007/978-1-61779-501-5_2

32. Gaffoor, I., Brown, D. W., Plattner, R., Proctor, R. H., Qi, W., & Trail, F. (2005). Functional analysis of the polyketide synthase genes in the filamentous fungus Gibberella zeae (Anamorph Fusarium graminearum). Eukaryotic Cell, 4(11), 1926–1933. 10.1128/EC.4.11.1926-1933.2005/ASSET/8D076388-27A5-4F3F-821D-B2C2077FE928/ASSETS/GRAPHIC/ZEK0110525500002.JPEG

33. García-Estrada, C., Domínguez-Santos, R., Kosalková, K., & Martín, J. F. (2018). Transcription Factors Controlling Primary and Secondary Metabolism in Filamentous Fungi: The β-Lactam Paradigm. Fermentation 2018, Vol. 4, Page 47, 4(2), 47. 10.3390/FERMENTATION4020047

34. Gay, E. J., Soyer, J. L., Lapalu, N., Linglin, J., Fudal, I., Da Silva, C., Wincker, P., Aury, J. M., Cruaud, C., Levrel, A., Lemoine, J., Delourme, R., Rouxel, T., & Balesdent, M. H. (2021). Large-scale transcriptomics to dissect 2 years of the life of a fungal phytopathogen interacting with its host plant. BMC Biology, 19(1), 1–27. 10.1186/S12915-021-00989-3/FIGURES/8

35. Geissman, T. A., Verbiscar, A. J., Phinney, B. O., & Cragg, G. (1966). Studies on the biosynthesis of gibberellins from (−)-kaurenoic acid in cultures of Gibberella Fujikuroi. Phytochemistry, 5(5), 933–947. 10.1016/S0031-9422(00)82790-9

36. Giraud-Gatineau, A., Ayachit, G., Nieves, C., Dagbo, K. C., Bourhy, K., Pulido, F., Huete, S. G., Benaroudj, N., Picardeau, M., & Veyrier, F. J. (2024). Inter-species Transcriptomic Analysis Reveals a Constitutive Adaptation Against Oxidative Stress for the Highly Virulent Leptospira Species. Molecular Biology and Evolution, 41(4). 10.1093/molbev/msae066

37. Gomes, L. B., Ward, T. J., Badiale-Furlong, E., & Del Ponte, E. M. (2015). Species composition, toxigenic potential and pathogenicity of Fusarium graminearum species complex isolates from southern Brazilian rice. Plant Pathology, 64(4), 980–987. 10.1111/PPA.12332

38. Harris, L. J., Balcerzak, M., Johnston, A., Schneiderman, D., & Ouellet, T. (2016). Host-preferential Fusarium graminearum gene expression during infection of wheat, barley, and maize. Fungal Biology, 120(1), 111–123. 10.1016/J.FUNBIO.2015.10.010

39. Harrison, M. C., LaBella, A. L., Hittinger, C. T., & Rokas, A. (2022). The evolution of the GALactose utilization pathway in budding yeasts. Trends in Genetics, 38(1), 97–106. 10.1016/j.tig.2021.08.013

40. Hicks, C., Witte, T. E., Sproule, A., Hermans, A., Shields, S. W., Colquhoun, R., Blackman, C., Boddy, C. N., Subramaniam, R., & Overy, D. P. (2023). CRISPR-Cas9 Gene Editing and Secondary Metabolite Screening Confirm Fusarium graminearum C16 Biosynthetic Gene Cluster Products as Decalin-Containing Diterpenoid Pyrones. Journal of Fungi, 9. 10.3390/jof9070695

41. Hicks, J. K., Yu, J. H., Keller, N. P., & Adams, T. H. (1997). Aspergillus sporulation and mycotoxin production both require inactivation of the FadA Gα protein-dependent signaling pathway. The EMBO Journal, 16(16), 4916–4923. 10.1093/EMBOJ/16.16.4916

42. Janevska, S., Arndt, B., Niehaus, E. M., Burkhardt, I., Rösler, S. M., Brock, N. L., Humpf, H. U., Dickschat, J. S., & Tudzynski, B. (2016). Gibepyrone biosynthesis in the rice pathogen Fusarium fujikuroi is facilitated by a small polyketide synthase gene cluster. Journal of Biological Chemistry, 291, 27403–27420. 10.1074/jbc.M116.753053

43. Jia, L. J., Tang, H. Y., Wang, W. Q., Yuan, T. L., Wei, W. Q., Pang, B., Gong, X. M., Wang, S. F., Li, Y. J., Zhang, D., Liu, W., & Tang, W. H. (2019). A linear nonribosomal octapeptide from Fusarium graminearum facilitates cell-to-cell invasion of wheat. Nature Communications, 10(1). 10.1038/s41467-019-08726-9

44. Johns, L. E., Bebber, D. P., Gurr, S. J., & Brown, N. A. (2022). Emerging health threat and cost of Fusarium mycotoxins in European wheat. Nature Food 2022 3:12, 3(12), 1014–1019. 10.1038/s43016-022-00655-z

45. Jones, D. A. B., Rozano, L., Debler, J. W., Mancera, R. L., Moolhuijzen, P. M., & Hane, J. K. (2021). An automated and combinative method for the predictive ranking of candidate effector proteins of fungal plant pathogens. Scientific Reports 2021 11:1, 11(1), 1–13. 10.1038/s41598-021-99363-0

46. Katoh, K., Rozewicki, J., & Yamada, K. D. (2017). MAFFT online service: multiple sequence alignment, interactive sequence choice and visualization. Briefings in Bioinformatics. 10.1093/bib/bbx108

47. Kazan, K., & Gardiner, D. M. (2018). Transcriptomics of cereal–Fusarium graminearum interactions: what we have learned so far. In Molecular Plant Pathology (Vol. 19, Issue 3, pp. 764–778). Blackwell Publishing Ltd. 10.1111/mpp.12561

48. Keller, N. P. (2015). Translating biosynthetic gene clusters into fungal armor and weaponry. Nature Chemical Biology, 11(9), 671–677. 10.1038/nchembio.1897

49. Keller, N. P. (2019). Fungal secondary metabolism: regulation, function and drug discovery. Nature Reviews Microbiology, 17(3), 167–180. 10.1038/s41579-018-0121-1

50. Kessler, S. C., Zhang, X., McDonald, M. C., Gilchrist, C. L. M., Lin, Z., Rightmyer, A., Solomon, P. S., Gillian Turgeon, B., & Chooi, Y. H. (2020). Victorin, the host-selective cyclic peptide toxin from the oat pathogen Cochliobolus victoriae, is ribosomally encoded. Proceedings of the National Academy of Sciences of the United States of America, 117(39), 24243–24250. 10.1073/pnas.2010573117

51. Kim, D., Paggi, J. M., Park, C., Bennett, C., & Salzberg, S. L. (2019). Graph-based genome alignment and genotyping with HISAT2 and HISAT-genotype. Nature Biotechnology, 37(8), 907–915. 10.1038/s41587-019-0201-4

52. King, R., Urban, M., & Hammond-Kosack, K. E. (2017). Annotation of Fusarium graminearum (PH-1) version 5.0. Genome Announcements, 5(2). 10.1128/genomeA.01479-16

53. Klassen, J. L. (2010). Phylogenetic and Evolutionary Patterns in Microbial Carotenoid Biosynthesis Are Revealed by Comparative Genomics. PLOS ONE, 5(6), e11257. 10.1371/JOURNAL.PONE.0011257

54. Li, J., Cornelissen, B., & Rep, M. (2020). Host-specificity factors in plant pathogenic fungi. Fungal Genetics and Biology, 144, 103447. 10.1016/J.FGB.2020.103447

55. Machado, F. J., Kuhnem, P. R., Casa, R. T., McMaster, N., Schmale, D. G., Vaillancourt, L. J., & Del Ponte, E. M. (2021). The Dominance of Fusarium meridionale over F. graminearum Causing Gibberella Ear Rot in Brazil May Be Due to Increased Aggressiveness and Competitiveness. Phytopathology, 111(10), 1774–1781. 10.1094/PHYTO-11-20-0515-R/ASSET/IMAGES/LARGE/PHYTO-11-20-0515-RF3.JPEG

56. Malz, S., Grell, M. N., Thrane, C., Maier, F. J., Rosager, P., Felk, A., Albertsen, K. S., Salomon, S., Bohn, L., Schäfer, W., & Giese, H. (2005). Identification of a gene cluster responsible for the biosynthesis of aurofusarin in the Fusarium graminearum species complex. Fungal Genetics and Biology, 42(5), 420–433. 10.1016/J.FGB.2005.01.010

57. Maor, R., & Shirasu, K. (2005). The arms race continues: battle strategies between plants and fungal pathogens. Current Opinion in Microbiology, 8(4), 399–404. 10.1016/J.MIB.2005.06.008

58. Miethke, M., & Marahiel, M. A. (2007). Siderophore-Based Iron Acquisition and Pathogen Control. Microbiology and Molecular Biology Reviews, 71(3), 413–451. 10.1128/MMBR.00012-07/ASSET/F38EFE85-C452-4F56-829C-DE44DDD76BCE/ASSETS/GRAPHIC/ZMR0030721580010.JPEG

59. Miguel-Rojas, C., Cavinder, B., Townsend, J. P., & Trail, F. (2023). Comparative Transcriptomics of Fusarium graminearum and Magnaporthe oryzae Spore Germination Leading up To Infection. MBio, 14(1). 10.1128/MBIO.02442-22/SUPPL_FILE/MBIO.02442-22-S0010.XLSX

60. Mirocha, C. J., & Swanson, S. P. (1983). REGULATION OF PERITHECIA PRODUCTION IN FUSARIUM ROSEUM BY ZEARALENONE1. Journal of Food Safety, 5(1), 41–53. 10.1111/J.1745-4565.1983.TB00454.X

61. Moser Tralamazza, S., Nanchira Abraham, L., Reyes-Avila, C. S., Corrêa, B., & Croll, D. (2022). Histone H3K27 Methylation Perturbs Transcriptional Robustness and Underpins Dispensability of Highly Conserved Genes in Fungi. Molecular Biology and Evolution, 39(1). 10.1093/MOLBEV/MSAB323

62. Niehaus, E. M., Kim, H. K., Münsterkötter, M., Janevska, S., Arndt, B., Kalinina, S. A., Houterman, P. M., Ahn, I. P., Alberti, I., Tonti, S., Kim, D. W., Sieber, C. M. K., Humpf, H. U., Yun, S. H., Güldener, U., & Tudzynski, B. (2017). Comparative genomics of geographically distant Fusarium fujikuroi isolates revealed two distinct pathotypes correlating with secondary metabolite profiles. PLOS Pathogens, 13(10), e1006670. 10.1371/JOURNAL.PPAT.1006670

63. Nielsen, J. C., Prigent, S., Grijseels, S., Workman, M., Ji, B., & Nielsen, J. (2019). Comparative Transcriptome Analysis Shows Conserved Metabolic Regulation during Production of Secondary Metabolites in Filamentous Fungi. MSystems, 4(2). 10.1128/MSYSTEMS.00012-19/ASSET/D6BE2229-8A05-42D3-AD65-D2038941E4D0/ASSETS/GRAPHIC/MSYSTEMS.00012-19-F0005.JPEG

64. Nielsen, M. R., Wollenberg, R. D., Westphal, K. R., Sondergaard, T. E., Wimmer, R., Gardiner, D. M., & Sørensen, J. L. (2019). Heterologous expression of intact biosynthetic gene clusters in Fusarium graminearum. Fungal Genetics and Biology, 132, 103248. 10.1016/J.FGB.2019.103248

65. Nützmann, H.-W., Scazzocchio, C., & Osbourn, A. (2018). Metabolic Gene Clusters in Eukaryotes. Annual Review of Genetics, 52(1), annurev-genet-120417-031237. 10.1146/annurev-genet-120417-031237

66. Oide, S., Berthiller, F., Wiesenberger, G., Adam, G., & Turgeon, B. G. (2015). Individual and combined roles of malonichrome, ferricrocin, and TAFC siderophores in Fusarium graminearum pathogenic and sexual development. Frontiers in Microbiology, 5, 759. 10.3389/fmicb.2014.00759

67. Patron, N. J., Waller, R. F., Cozijnsen, A. J., Straney, D. C., Gardiner, D. M., Nierman, W. C., & Howlett, B. J. (2007). Origin and distribution of epipolythiodioxopiperazine (ETP) gene clusters in filamentous ascomycetes. BMC Evolutionary Biology, 7(1), 174. 10.1186/1471-2148-7-174

68. Peregrín-Alvarez, J. M., Sanford, C., & Parkinson, J. (2009). The conservation and evolutionary modularity of metabolism. Genome Biology, 10(6), 1–17. 10.1186/GB-2009-10-6-R63/FIGURES/6

69. Pontecorvo, G., Roper, J. A., Chemmons, L. M., Macdonald, K. D., & Bufton, A. W. J. (1953). The genetics of Aspergillus nidulans. Advances in Genetics, 5(C), 141–238. 10.1016/S0065-2660(08)60408-3

70. Proctor, R. H., McCormick, S. P., Kim, H.-S., Cardoza, R. E., Stanley, A. M., Lindo, L., Kelly, A., Brown, D. W., Lee, T., Vaughan, M. M., Alexander, N. J., Busman, M., & Gutiérrez, S. (2018). Evolution of structural diversity of trichothecenes, a family of toxins produced by plant pathogenic and entomopathogenic fungi. PLOS Pathogens, 14(4), e1006946. 10.1371/journal.ppat.1006946

71. Rambaut A. (2009). Figtree.Tree figure drawing tool.

72. Rank, C., Nielsen, K. F., Larsen, T. O., Varga, J., Samson, R. A., & Frisvad, J. C. (2011). Distribution of sterigmatocystin in filamentous fungi. Fungal Biology, 115(4–5), 406–420. 10.1016/J.FUNBIO.2011.02.013

73. Ritchie, M. E., Phipson, B., Wu, D., Hu, Y., Law, C. W., Shi, W., & Smyth, G. K. (2015). limma powers differential expression analyses for RNA-sequencing and microarray studies. Nucleic Acids Research, 43(7), e47–e47. 10.1093/NAR/GKV007

74. Robinson, M. D., McCarthy, D. J., & Smyth, G. K. (2010). edgeR: a Bioconductor package for differential expression analysis of digital gene expression data. Bioinformatics, 26(1), 139–140. 10.1093/bioinformatics/btp616

75. Rokas, A., Mead, M. E., Steenwyk, J. L., Raja, H. A., & Oberlies, N. H. (2020). Biosynthetic gene clusters and the evolution of fungal chemodiversity. In Natural Product Reports (Vol. 37, Issue 7, pp. 868–878). Royal Society of Chemistry. 10.1039/c9np00045c

76. Rokas, A., Wisecaver, J. H., & Lind, A. L. (2018). The birth, evolution and death of metabolic gene clusters in fungi. Nature Reviews Microbiology, 1. 10.1038/s41579-018-0075-3

77. Severinsen, M. M., Westphal, K. R., Terp, M., Sørensen, T., Olsen, A., Bachleitner, S., Studt-Reinhold, L., Wimmer, R., Sondergaard, T. E., & Sørensen, J. L. (2023). Filling out the gaps – identification of fugralins as products of the PKS2 cluster in Fusarium graminearum. Frontiers in Fungal Biology, 4. 10.3389/ffunb.2023.1264366

78. Shimizu, K., & Keller, N. P. (2001). Genetic Involvement of a cAMP-Dependent Protein Kinase in a G Protein Signaling Pathway Regulating Morphological and Chemical Transitions in Aspergillus nidulans. Genetics, 157(2), 591–600. 10.1093/GENETICS/157.2.591

79. Sieber, C. M. K., Lee, W., Wong, P., Münsterkötter, M., Mewes, H.-W., Schmeitzl, C., Varga, E., Berthiller, F., Adam, G., & Güldener, U. (2014). The Fusarium graminearum Genome Reveals More Secondary Metabolite Gene Clusters and Hints of Horizontal Gene Transfer. PLoS ONE, 9(10), e110311. 10.1371/journal.pone.0110311

80. Somu, R. V., Boshoff, H., Qiao, C., Bennett, E. M., Barry, C. E., & Aldrich, C. C. (2006). Rationally-designed nucleoside antibiotics that inhibit siderophore biosynthesis of Mycobacterium tuberculosis. Journal of Medicinal Chemistry, 49(1), 31–34. 10.1021/JM051060O/SUPPL_FILE/JM051060OSI20051122_101858.PDF

81. Sondergaard, T. E., Hansen, F. T., Purup, S., Nielsen, A. K., Bonefeld-Jørgensen, E. C., Giese, H., & Sørensen, J. L. (2011). Fusarin C acts like an estrogenic agonist and stimulates breast cancer cells in vitro. Toxicology Letters, 205(2), 116–121. 10.1016/J.TOXLET.2011.05.1029

82. Stamatakis, A. (2014). RAxML version 8: a tool for phylogenetic analysis and post-analysis of large phylogenies. Bioinformatics, 30(9), 1312–1313. 10.1093/bioinformatics/btu033

83. Stanke, M., & Morgenstern, B. (2005). AUGUSTUS: a web server for gene prediction in eukaryotes that allows user-defined constraints. Nucleic Acids Research, 33(Web Server), W465–W467. 10.1093/nar/gki458

84. Stephens, A. E., Gardiner, D. M., White, R. G., Munn, A. L., & Manners, J. M. (2008). Phases of Infection and Gene Expression of Fusarium graminearum During Crown Rot Disease of Wheat. 10.1094/MPMI-21-12-1571, 21(12), 1571–1581. https://doi.org/10.1094/MPMI-21-12-1571

85. Studt, L., Strauss, J., Janevska, S., Arndt, B., Boedi, S., Sulyok, M., Humpf, H.-U., & Tudzynski, B. (2017). Lack of the COMPASS Component Ccl1 Reduces H3K4 Trimethylation Levels and Affects Transcription of Secondary Metabolite Genes in Two Plant–Pathogenic Fusarium Species. Frontiers in Microbiology, 7, 229533. 10.3389/FMICB.2016.02144

86. Takahashi, H., Umemura, M., Ninomiya, A., Kusuya, Y., Shimizu, M., Urayama, S. I., Watanabe, A., Kamei, K., Yaguchi, T., & Hagiwara, D. (2021). Interspecies Genomic Variation and Transcriptional Activeness of Secondary Metabolism-Related Genes in Aspergillus Section Fumigati. Frontiers in Fungal Biology, 2, 656751. 10.3389/FFUNB.2021.656751/BIBTEX

87. Tralamazza, S. M., Rocha, L. O., Oggenfuss, U., Corrêa, B., Croll, D., & Rose, L. (2019). Complex Evolutionary Origins of Specialized Metabolite Gene Cluster Diversity among the Plant Pathogenic Fungi of the Fusarium graminearum Species Complex. Genome Biology and Evolution, 11(11), 3106–3122. 10.1093/gbe/evz225

88. Tsai, H. F., Chang, Y. C., Washburn, R. G., Wheeler, M. H., & Kwon-Chung, K. J. (1998). The Developmentally Regulated alb1 Gene of Aspergillus fumigatus: Its Role in Modulation of Conidial Morphology and Virulence. Journal of Bacteriology, 180(12), 3031. 10.1128/JB.180.12.3031-3038.1998

89. Tu, Q., Wang, L., An, Q., Shuai, J., Xia, X., Dong, Y., Zhang, X., Li, G., & He, Y. (2023). Comparative transcriptomics identifies the key in planta-expressed genes of Fusarium graminearum during infection of wheat varieties. Frontiers in Genetics, 14, 1166832. 10.3389/FGENE.2023.1166832/BIBTEX

90. van der Lee, T., Zhang, H., van Diepeningen, A., & Waalwijk, C. (2015). Biogeography of *Fusarium graminearum* species complex and chemotypes: a review. Food Additives & Contaminants: Part A, 32(4), 453–460. 10.1080/19440049.2014.984244

91. Vogt, E., & Künzler, M. (2019). Discovery of novel fungal RiPP biosynthetic pathways and their application for the development of peptide therapeutics. Applied Microbiology and Biotechnology, 1–15. 10.1007/s00253-019-09893-x

92. Walter, S., Nicholson, P., & Doohan, F. M. (2010). Action and reaction of host and pathogen during Fusarium head blight disease. New Phytologist, 185(1), 54–66. 10.1111/J.1469-8137.2009.03041.X

93. Wang, J. H., Ndoye, M., Zhang, J. B., Li, H. P., & Liao, Y. C. (2011). Population Structure and Genetic Diversity of the Fusarium graminearum Species Complex. Toxins 2011, Vol. 3, Pages 1020-1037, 3(8), 1020–1037. 10.3390/TOXINS3081020

94. Wilkinson, H. H., Ramaswamy, A., Sung, C. S., & Keller, N. P. (2004). Increased conidiation associated with progression along the sterigmatocystin biosynthetic pathway. Mycologia, 96(6), 1190–1198. 10.1080/15572536.2005.11832867

95. Woelflingseder, L., Warth, B., Vierheilig, I., Schwartz-Zimmermann, H., Hametner, C., Nagl, V., Novak, B., Šarkanj, B., Berthiller, F., Adam, G., & Marko, D. (2019). The Fusarium metabolite culmorin suppresses the in vitro glucuronidation of deoxynivalenol. Archives of Toxicology, 93, 1729–1743. 10.1007/s00204-019-02459-w

96. Wolf, J. C., & Mirocha, C. J. (2011). Regulation of sexual reproduction in Gibberella zeae (Fusarium roseum ’Graminearum’) by F-2 (Zearalenone). 10.1139/M73-117, 19(6), 725–734. https://doi.org/10.1139/M73-117

97. Won, T. H., Bok, J. W., Nadig, N., Venkatesh, N., Nickles, G., Greco, C., Lim, F. Y., González, J. B., Turgeon, B. G., Keller, N. P., & Schroeder, F. C. (2022). Copper starvation induces antimicrobial isocyanide integrated into two distinct biosynthetic pathways in fungi. Nature Communications 2022 13:1, 13(1), 1–14. 10.1038/s41467-022-32394-x

98. Yamada, A., Kataoka, T., & Nagai, K. (2000). The fungal metabolite gliotoxin: immunosuppressive activity on CTL-mediated cytotoxicity. Immunology Letters, 71(1), 27–32. 10.1016/S0165-2478(99)00155-8

99. Yang, Y., Li, M. X., Duan, Y. B., Li, T., Shi, Y. Y., Zhao, D. L., Zhou, Z. H., Xin, W. J., Wu, J., Pan, X. Y., Li, Y. J., Zhu, Y. Y., & Zhou, M. G. (2018). A new point mutation in β2-tubulin confers resistance to carbendazim in Fusarium asiaticum. Pesticide Biochemistry and Physiology, 145, 15–21. 10.1016/J.PESTBP.2017.12.006

100. Zhang, C., Sayyari, E., & Mirarab, S. (2017). ASTRAL-III: Increased Scalability and Impacts of Contracting Low Support Branches (pp. 53–75). Springer, Cham. 10.1007/978-3-319-67979-2_4

101. Zhang, H., van der Lee, T., Waalwijk, C., Chen, W., Xu, J., Xu, J., Zhang, Y., & Feng, J. (2012). Population Analysis of the Fusarium graminearum Species Complex from Wheat in China Show a Shift to More Aggressive Isolates. PLOS ONE, 7(2), e31722. 10.1371/JOURNAL.PONE.0031722

102. Zhang, X.-W., Jia, L.-J., Zhang, Y., Jiang, G., Li, X., Zhang, D., & Tang, W.-H. (2012). In Planta Stage-Specific Fungal Gene Profiling Elucidates the Molecular Strategies of Fusarium graminearum Growing inside Wheat Coleoptiles. The Plant Cell, 24(12), 5159–5176. 10.1105/tpc.112.105957

103. Zhao, S., Zhang, K., Lin, C., Cheng, M., Song, J., Ru, X., Wang, Z., Wang, W., & Yang, Q. (2022). Identification of a Novel Pleiotropic Transcriptional Regulator Involved in Sporulation and Secondary Metabolism Production in Chaetomium globosum. International Journal of Molecular Sciences, 23(23), 14849. 10.3390/IJMS232314849/S1

