## Supplementary Figures for "Wheat infection by *Fusarium graminearum* species complex members is facilitated by a transcriptionally conserved non-ribosomal peptide synthetase gene cluster"

### Supplementary Information

Tralamazza *et al.*

**Supplementary Figures S1-6**

**Supplementary Tables S1-11** (see separate file)

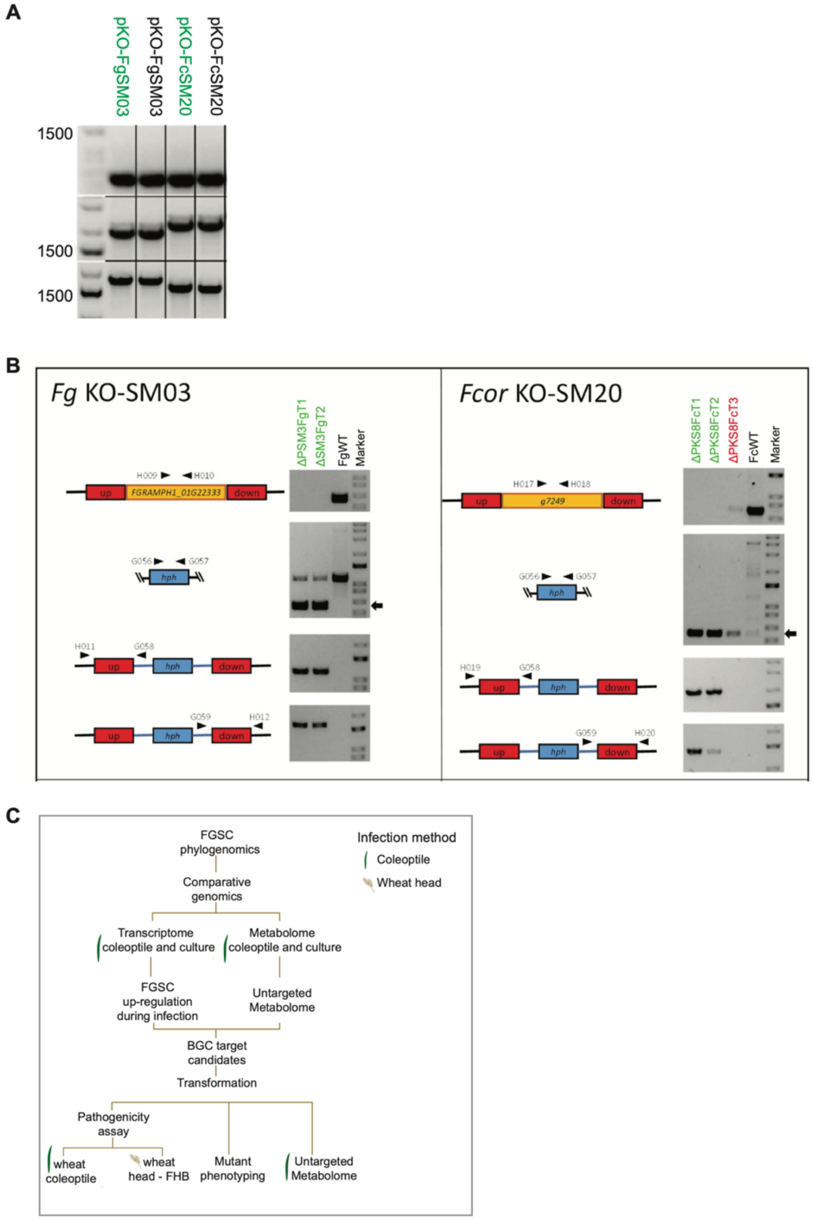


**Supplementary Figure S1.** Biosynthetic gene cluster deletion mutants. A) Plasmid construct validation with PCR. Green color highlight PCR and sanger sequencing positive results selected for downstream applications. B) PCR validation of fungal knock-out transformants. Green color refers to validated targets. C) Schematics describing the current study.

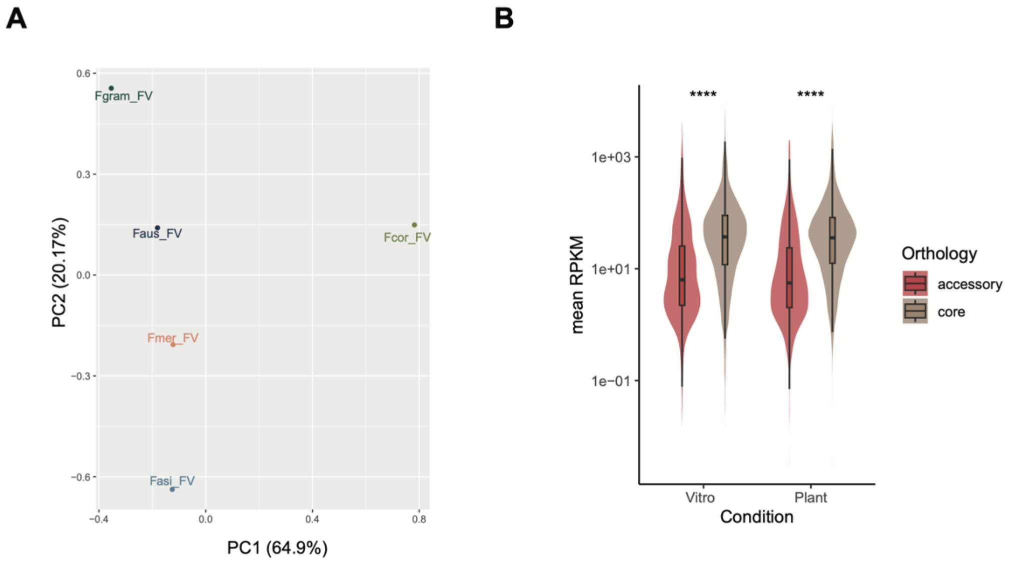


**Supplementary Figure S2.** A) Transcriptomic profile of FGSC species during *in vitro* culture based on principal components (PCs) analysis PC1 and PC2. Colors refer to the species (Fgram: *F. graminearum s.s,* Fmer: *F. meridionale,* Fcor: *F. cortaderiae,* Faus: *F. austroamericanum* and Fasi: *F. asiaticum*). B) FGSC mean gene expression difference between core and accessory genes. Wilcoxon tests with adjusted P-values (Holm method). Ns: p> 0.05.

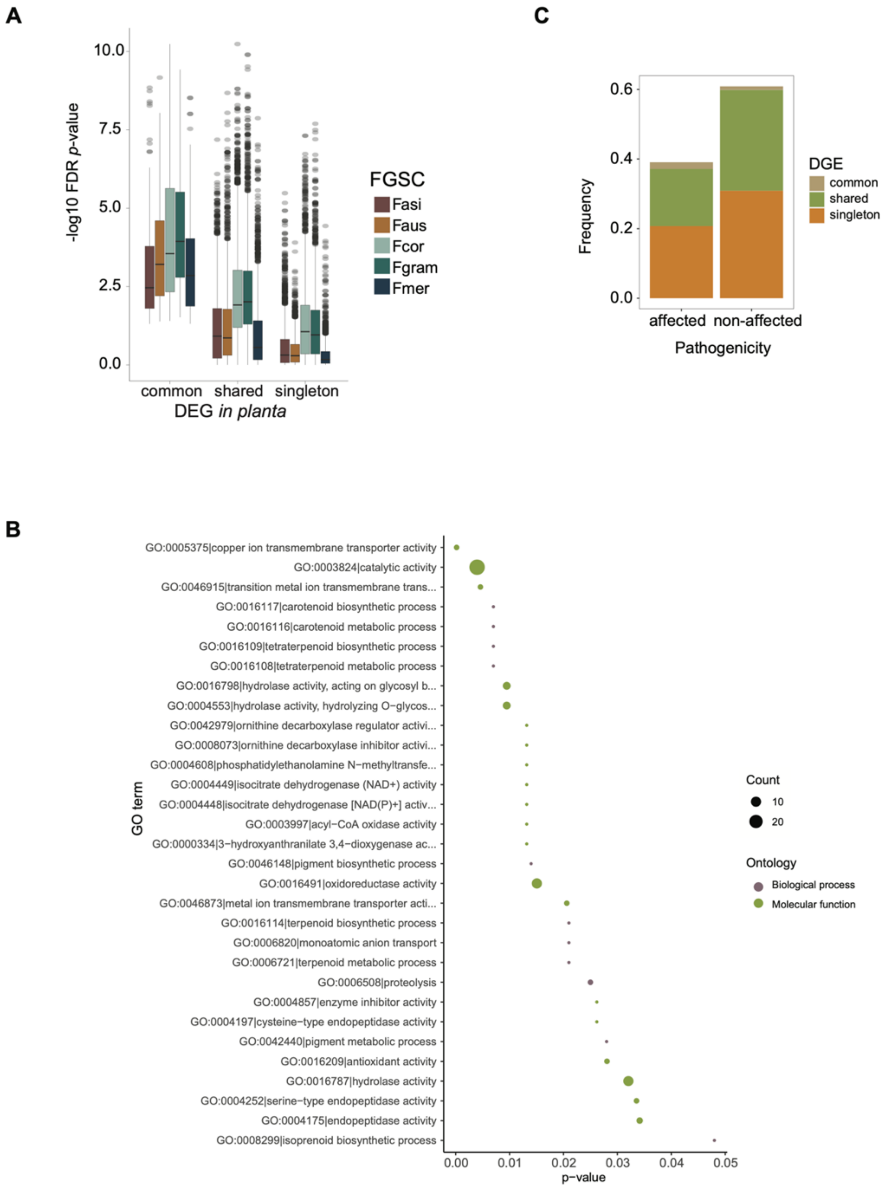


**Supplementary Figure S3.** A) DGE significance (-logFDR) distribution based on the DGE categories. Colors indicate DGE genes shared between FGSC species. Common category refers to DGE in all species, shared refers to less than five species and singleton to a single species. Unchanged are genes not differentially expressed between conditions. B) GO term enrichment analysis of DGE common genes. Circle size represents the enriched GO term size. Fisher test was performed. C) Frequency of DGE with publicly available mutant screens for effects on host infection based on the curated PHI-base. Pathogenicity affected phenotypes include increased/reduced virulence, loss of pathogenicity, and lethality.

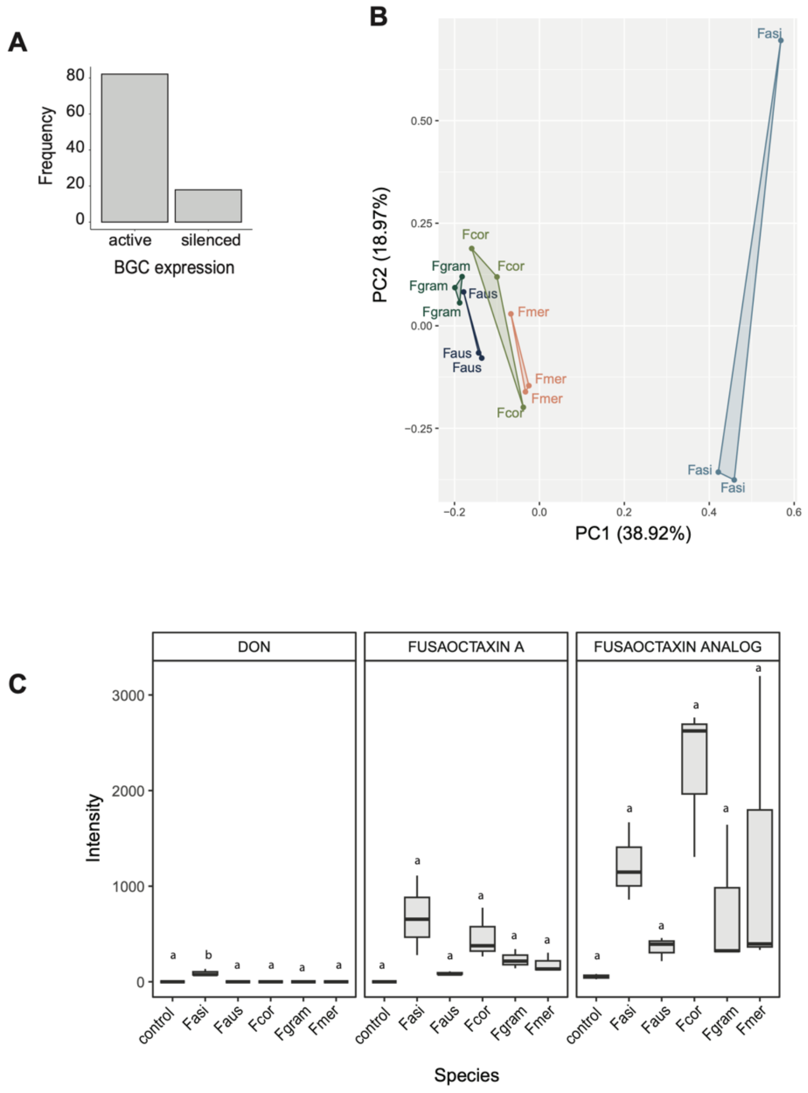


**Supplementary Figure S4.** A) Frequency of active and silenced BGCs under the tested conditions. B) Metabolomic profile of FGSC species during *in vitro* culture based on principal components (PCs) analysis PC1 and PC2. Colors refer to the species (Fgram: *F. graminearum s.s,* Fmer: *F. meridionale,* Fcor: *F. cortaderiae,* Faus: *F. austroamericanum* and Fasi: *F. asiaticum*). C) Metabolite intensity of deoxynivalenol, Fusaoctaxin A and analog across FGSC species during *in vitro* culture. Different letters above the boxplot identify significantly different groups according to ANOVA and Tukey test.

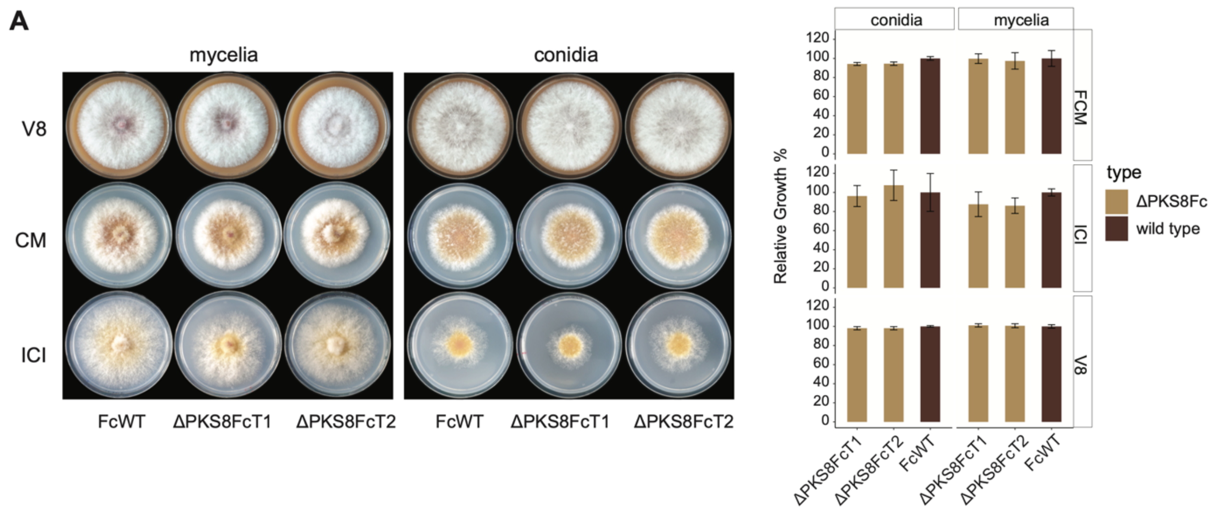


**Supplementary Figure S5.** Representative images of 5-day-old cultures *F. cortaderiae* wild strain (FcWT) and PKS8 deleted mutants (ΔPKS8Fc). Right plot refers to the relative growth of each condition tested.

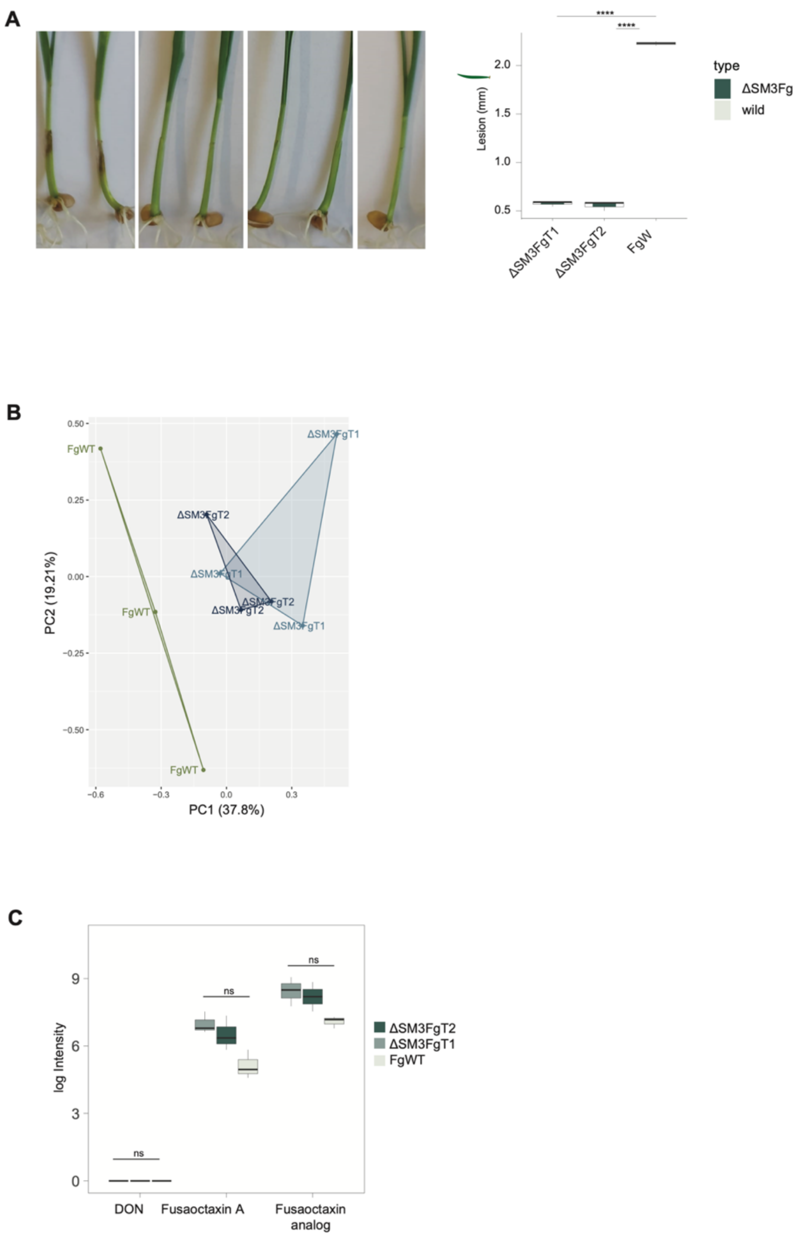


**Supplementary Figure S6.** A) Representative images of wheat seedlings at 4 days after coleoptile inoculation with *F. graminearum* wild strain (FgWT) and sm3 deleted mutants (ΔSM3Fg). Control was inoculated with water. Right boxplot refers to the distribution of lesion size in the tested strains. B) Metabolomic profile of wild (FgWT) and mutant strains (ΔSM3Fg) during *in vitro* culture based on principal components (PCs) analysis PC1 and PC2. Colors refer to the strains. C) Metabolite intensity of deoxynivalenol, Fusaoctaxin A and analog in the wild (FgWT) and mutant strains (ΔSM3Fg) during *in vitro* culture. Wilcoxon tests with adjusted P-values (Holm method). Ns: $p > 0.05$.
